# *Cul3* regulates cytoskeleton protein homeostasis and cell migration during a critical window of brain development

**DOI:** 10.1101/2020.01.10.902064

**Authors:** Jasmin Morandell, Lena A. Schwarz, Bernadette Basilico, Saren Tasciyan, Armel Nicolas, Christoph Sommer, Caroline Kreuzinger, Christoph P. Dotter, Lisa S. Knaus, Zoe Dobler, Emanuele Cacci, Johann G. Danzl, Gaia Novarino

**Affiliations:** Institute of Science and Technology (IST) Austria, Klosterneuburg, Austria; Department of Biology and Biotechnology “Charles Darwin”, Sapienza, University of Rome

## Abstract

*De novo* loss of function mutations in the ubiquitin ligase-encoding gene *Cullin3* (*CUL3)* lead to autism spectrum disorder (ASD). Here, we used *Cul3* mouse models to evaluate the consequences of *Cul3* mutations *in vivo.* Our results show that *Cul3* haploinsufficient mice exhibit deficits in motor coordination as well as ASD-relevant social and cognitive impairments. *Cul3* mutant brain displays cortical lamination abnormalities due to defective neuronal migration and reduced numbers of excitatory and inhibitory neurons. In line with the observed abnormal columnar organization, *Cul3* haploinsufficiency is associated with decreased spontaneous excitatory and inhibitory activity in the cortex. At the molecular level, employing a quantitative proteomic approach, we show that *Cul3* regulates cytoskeletal and adhesion protein abundance in mouse embryos. Abnormal regulation of cytoskeletal proteins in *Cul3* mutant neuronal cells results in atypical organization of the actin mesh at the cell leading edge, likely causing the observed migration deficits. In contrast to these important functions early in development, *Cul3* deficiency appears less relevant at adult stages. In fact, induction of *Cul3* haploinsufficiency in adult mice does not result in the behavioral defects observed in constitutive *Cul3* haploinsufficient animals. Taken together, our data indicate that *Cul3* has a critical role in the regulation of cytoskeletal proteins and neuronal migration and that ASD-associated defects and behavioral abnormalities are primarily due to *Cul3* functions at early developmental stages.

## Main

The past decade has seen a major effort to elucidate the genetic underpinnings of autism spectrum disorders (ASDs). Whole exome sequencing of large patient cohorts and their unaffected family members has identified hundreds of ASD risk loci^1-5^. However, the molecular and cellular functions of the majority of the identified genes remain poorly understood. One of the identified high-risk ASD genes encodes the E3 ubiquitin ligase Cullin 3 (Cul3)^1-3,6-10^.

E3 ubiquitin ligases regulate cellular protein composition by providing target recognition and specificity to the ubiquitin-dependent proteasomal degradation pathway^11^. CUL3 is a conserved protein of the Cullin family, comprising eight members, which contain a conserved cullin homology domain, named after its ability to select cellular proteins for degradation. *CUL3* ASD-associated genetic variants are most often *de novo* missense or loss of function (loF) mutations, dispersed throughout the entire gene and affecting distinct protein domains. In addition to the ASD core symptoms, patients with *CUL3 de novo* loF mutations can present with several comorbidities including varying levels of intellectual disability (ID), attention deficit hyperactivity disorder (ADHD), sleep disturbances, motor deficits and facial dysmorphic features^10,12,13^. The only known exception is the deletion of *CUL3* exon 9 by a specific dominant splice site variant causing a severe form of pseudohypoaldosteronism type II (PHAII), featuring hypertension, hyperkalemia, and metabolic acidosis but not ASD^14,15,16^. Despite the well-understood process of CUL3-mediated protein ubiquitination and degradation^11^, its target proteins in the developing central nervous system and its role in brain development and adult brain functions remain largely unknown.

In mouse, complete deletion of *Cul3* is lethal at embryonic day (E) 7.5 due to gastrulation defects and aberrant cell cycle regulation caused by accumulation of Cyclin E, a regulator of mitotic S-phase^17^. Here, we employed heterozygous *Cul3* knockout mouse lines to model the pathophysiological consequences of *CUL3* mutations. We show that *Cul3* is crucial for the development of the central nervous system and its haploinsufficiency leads to behavioral abnormalities including altered social preference, sensory hyper-reactivity, motor coordination defects and cognitive impairment. Analysis of the cerebral cortex architecture links *Cul3* haploinsufficiency to lamination abnormalities, which correlate with cortical network activity defects. Concordantly, forebrain-specific homozygous deletion of *Cul3* leads to severe cortical malformations in newborn animals. Further *in vivo* and *in vitro* experiments underscored Cul3’s importance for neural cell migration by substantiating its role in regulating cytoskeletal and adhesion protein levels, and actin cytoskeleton dynamics. Importantly, deletion of *Cul3* in adult animals does not lead to behavioral abnormalities, suggesting that *Cul3* is critical during early developmental stages. Altogether, our results highlight a pivotal role for *Cul3* in normal brain development and suggest the existence of a critical temporal window for the treatment of *CUL3*-linked ASD.

## Results

### Behavioral defects in *Cul3* haploinsufficient animals

To model ASD-linked mutations, we studied a constitutive heterozygous *Cul3* knockout (*Cul3*^*+/-*^) mouse^18^ (Supplementary Fig. 1a). As predicted, *Cul3*^*+/-*^ animals show a significant decrease in Cul3 protein in the brain, to approximately 50% of wild-type levels (Supplementary Fig. 1b, Supplementary Table 1). Importantly, Cul3 protein reduction is equal in all brain areas tested (Supplementary Fig. 1b), thus resembling patients with germline mutations.

**Figure 1.**
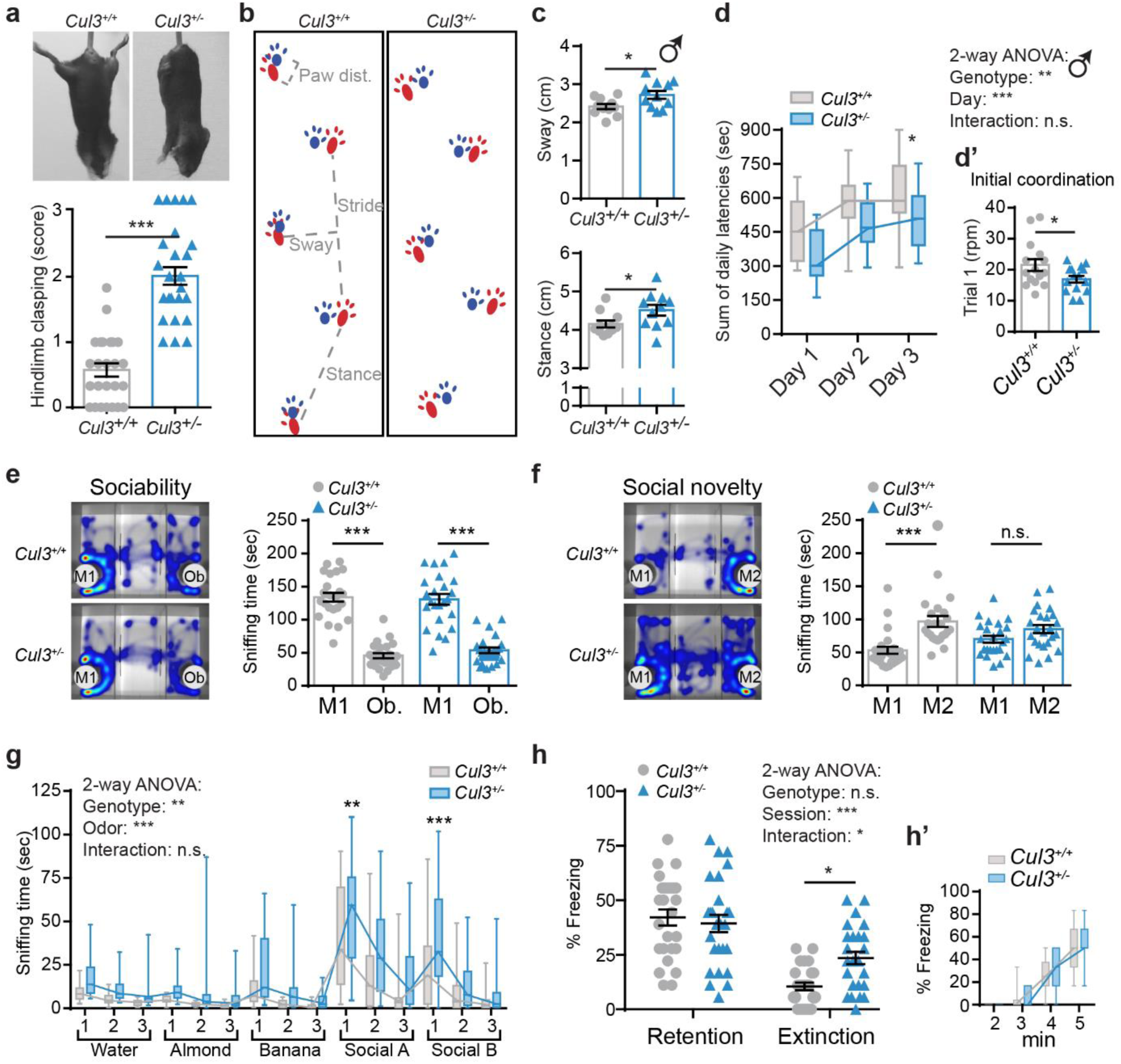
Behavioral defects in *Cul3* haploinsufficient mice. **a**, Representative images of hindlimb clasping in adult *Cul3*^*+/-*^ mice, not observed in their wild-type littermate controls (a, top) and scoring from 0-1 (normal) to 3 (most severe) (a, bottom, *n*= 25 animals, females (*n*= 11) and males (*n*= 14), per genotype; ***P<0.001; two-tailed Mann-Whitney U-test). **b**, Representative images of *Cul3*^*+/+*^ and *Cul3*^*+/-*^ strides, forepaws in blue and hindpaws in red. **c**, Altered gait of the *Cul3* haploinsufficient male mice evidenced by inter-genotype comparison of sway (c, top) and stance length (c, bottom) (*n*= 11 male mice per genotype, littermates; *P<0.05; two-tailed Mann-Whitney U test or two-tailed t-test). **d**-**d’**, Accelerating RotaRod test revealing defects in motor learning and coordination in *Cul3*^*+/-*^ mice. Shown are the sum of daily latencies of three trials per day on three consecutive days (d) and the final rpm on day one - trial 1, as measure of initial coordination (d’) (*n*= 15 male mice per genotype; *P<0.05; 2-way ANOVA and Sidak’s multiple comparison test and unpaired two-tailed t-test). Data for female animals in Supplementary Figure 2. **e**-**f**, Representative heat maps of the three-chamber social interaction test (left) and quantification of interaction times (right). Sociability: *Cul3*^*+/-*^ and control mice spend more time with a stranger mouse (M1) rather than an object (Ob.) (e); Social novelty: *Cul3*^*+/-*^ mice do not prefer a novel stranger (M2) over the already familiar mouse (M1) (f) (*n*= 24 mice, females (*n*= 11) and males (*n*= 13), per genotype; ***P<0.001, n.s. not significant; 1-way ANOVA and Sidak’s multiple comparison test). **g**, Both genotypes distinguish and familiarize to non-social and social odors in the olfaction habituation and dishabituation test, yet *Cul3*^*+/-*^ mutant mice are hyper-reactive to the presentation of social odors (*n*= 24 mice, females (*n*= 11) and males (*n*= 13), per genotype; **P<0.01, ***P<0.001, n.s. not significant; 2-way ANOVA and Sidak’s multiple comparison test; details in Supplementary Tables 1,2). **h**-**h’**, Contextual fear-conditioned memory retention and extinction scored as percent freezing during a 3 min exposure to the context (h), and fear acquisition training (h’) (*n*= 26 mice, females (*n*= 7) and males (*n*= 19), per genotype; *P<0.05, ***P<0.001, n.s., not significant; 2-way ANOVA interaction: (F1,100)= 6.18 p= 0.015; Sidak’s multiple comparisons test: Extinction p= 0.027). Data presented either as mean ± SEM, as well as scatter plot (a,c,d’,e,f,h) or as box and whiskers, min. to max., (d,g,h’). Detailed statistics are provided in Supplementary Table 1.

Although *Cul3* haploinsufficient mice have a slightly reduced body weight at birth, their weight is comparable to control animals as adults (Supplementary Fig. 1c), while the brain to body weight ratio is unaffected in mutant newborn and adult mice (Supplementary Fig. 1d). Adult *Cul3*^*+/-*^ mice present with hindlimb clasping (Fig. 1a) and mild gait abnormalities, such as increased sway and stance length (Fig. 1b,c; Supplementary Fig. 2a,b), phenotypes which are observed in other ASD mouse models^19,20^ and indicative of cerebellar dysfunctions^21^. Further indicating motor defects, *Cul3*^*+/-*^ mice underperform when challenged on the accelerating RotaRod (Fig. 1d,d’; Supplementary Fig. 2c,c’), a task requiring formation and consolidation of a repetitive motor routine^22,23^. Male, but not female, *Cul3* haploinsufficient mice show reduced initial coordination compared to their wild-type littermates (Fig. 1d’; Supplementary Fig. 2c’). In addition, mutant mice of both sexes do not reach the same level of motor performance as their healthy counterparts by the end of the third day of trials, suggesting motor learning impairments (Fig. 1d, Supplementary Fig. 1c). Motor defects of *Cul3*^*+/-*^ mice, however, do not affect exploratory behavior in the open field (Supplementary Fig. 2d), nor on the elevated plus maze, where *Cul3*^*+/-*^ animals do not show differences in anxiety-like behaviors (Supplementary Fig. 2e).

**Figure 2.**
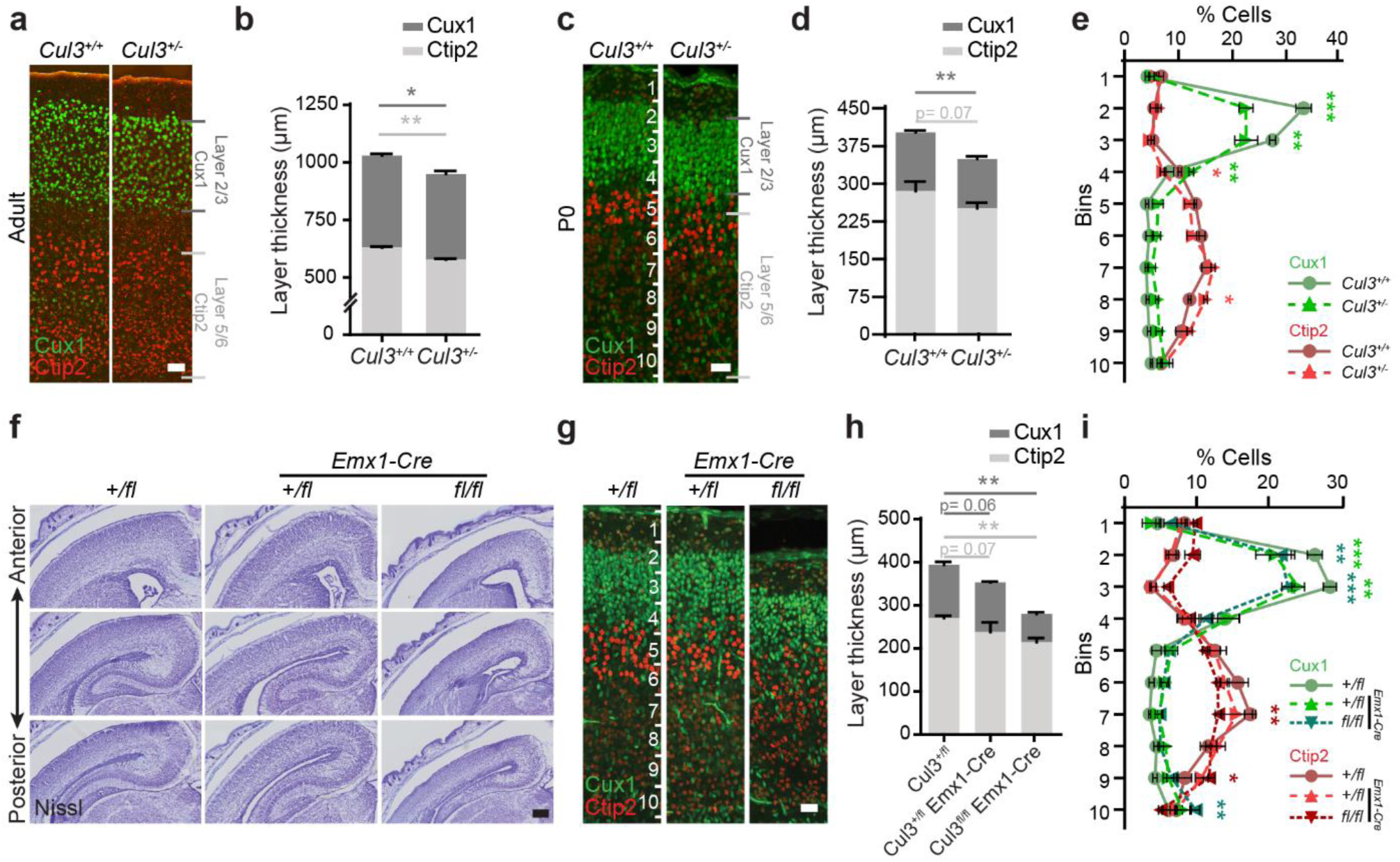
Abnormal lamination of the somatosensory cortex in *Cul3* mutant mice. **a**-**d**, Immunofluorescent stainings for Ctip2 and Cux1 on coronal brain sections revealed laminar thinning in adults (a,b) and newborn (P0) *Cul3*^*+/-*^ animals (c,d) (*n*(adults)= 3 littermates per genotype; *n*(P0)= 6 littermates per genotype; *P<0.05; 2-way ANOVA and Sidak’s multiple comparison test). **e**, Bin-wise comparison of relative cell numbers (in %) revealed a shifted Cux1/Ctip2 layer profile, indicating laminar defects at P0 (*n*= 3 littermates per genotype; *P<0.05, **P<0.01, ***P<0.001; 2-way ANOVA, Sidak’s multiple comparison test). **f**, Nissl-staining of P0 coronal, *Cul3*^*+/fl*^, *Cul3*^*+/fl*^ *Emx1-Cre and Cul3*^*fl/fl*^ *Emx1-Cre* brain sections show severe brain malformations in *Cul3*^*fl/fl*^ *Emx1-Cre* pups. **g**-**i**, Immunofluorescent stainings with antibodies against Ctip2 and Cux1 reveal cortical laminar thinning in both *Cul3*^*+/fl*^ *Emx1-Cre and Cul3*^*fl/fl*^ *Emx1-Cre* pups and bin-wise comparison of relative cell numbers show a shifted Cux1/Ctip2 layer profile at P0 (*n*= 3 littermates per genotype; *P<0.05, **P<0.01, ***P<0.001; 2-way ANOVA, Sidak’s multiple comparison test). Data presented as stacked bar-plots of mean ± SEM in (b,d,h) and connected mean ± SEM in (e) and (i). Scale bars: 50 µm in (a), 25 µm in (c,g) and 200 µm in (f); numbers in (c,g) indicate depth in the cortex. Detailed statistics are provided in Supplementary Table 1.

Next, we subjected *Cul3*^*+/-*^ animals to classical sociability tests. In the three-chamber test, similarly to wild types, *Cul3*^*+/-*^ mice show preference for a mouse (M1) over an object (Ob.) (Fig. 1e). However, in the second phase of the assay, mutant mice show no preference for a stranger mouse (M2) over a familiar animal (M1), a preference displayed by control animals (Fig. 1f). We thus concluded that haploinsufficiency of the *Cul3* gene is associated with reduced social memory. As social recognition is mainly achieved via olfaction in rodents^24,25^, we assessed the ability of mutant animals to distinguish and familiarize with non-social and social odors. In the odor discrimination and habituation test (ODHD)^26^, both wild-type and *Cul3*^*+/-*^ animals successfully recognize newly and already presented odors (Supplementary Table 2). However, mutant mice spend significantly more time exploring odor-embedded cotton swabs, and are hyper-reactive to the presentation of social odors (2-way ANOVA: genotype (F1,46)= 10.07, p= 0.003) (Fig. 1g, Supplementary Table 1,2). Thus, despite mutant animals spending significantly more time sniffing social odors than controls, they are able to distinguish between two different social odors, indicating that defects in social memory are not directly related to odor discrimination issues. Finally, we employed a well-established memory test^27^ to assess how *Cul3* haploinsufficiency affects learning. Contextual fear conditioning revealed normal fear acquisition and memory retention in *Cul3*^*+/-*^ mice. However, mutant animals exhibit reduced ability to extinguish the aversive memory after extinction training, pointing towards abnormal cognition (Fig. 1h,h’).

In summary, our analysis indicates that *Cul3* haploinsufficiency leads to abnormalities in several behavioral paradigms, potentially associated with dysfunction of different brain areas and/or dysfunctional brain connectivity.

### *Cul3* haploinsufficiency is associated with abnormal brain development

Temporally, in the mouse brain, *Cul3* expression peaks at E14.5 and E16.5 (Supplementary Fig. 3a,b). Spatially, it is predominantly expressed in the cortex and hippocampus (Supplementary Fig. 3c), in glutamatergic and inhibitory neurons (Supplementary Fig. 3d). These data are in line with the *CUL3* expression profile in the human brain (Supplementary Fig. 3e-g) and point towards an important role for *Cul3* in neuronal cells during early brain development. Thus, to understand whether behavioral defects are accompanied by neuroanatomical changes, we performed crystal violet (Nissl) stainings of adult brain sections obtained from *Cul3* mutant and wild-type mice (Supplementary Fig. 4). Gross brain morphology appears normal but we observed a slight reduction in cortical thickness and cerebellar area (Supplementary Fig. 4e’’,e’’’). To investigate this neuroanatomical phenotype more closely, we stained the somatosensory cortex for Cux1 (upper layers 2/3) and Ctip2 (lower layers 5/6) (Fig. 2a-d). Quantifications in adult mice revealed that the *Cul3* heterozygous mutation results in a mild decrease in upper and lower cortical layer thickness (Fig. 2b), a defect present already at birth (Fig. 2d). In addition, we found that the distribution of Cux1+ and Ctip2+ cells is shifted toward lower cortical locations, indicative of abnormal cortical lamination (Fig. 2e).

**Figure 3.**
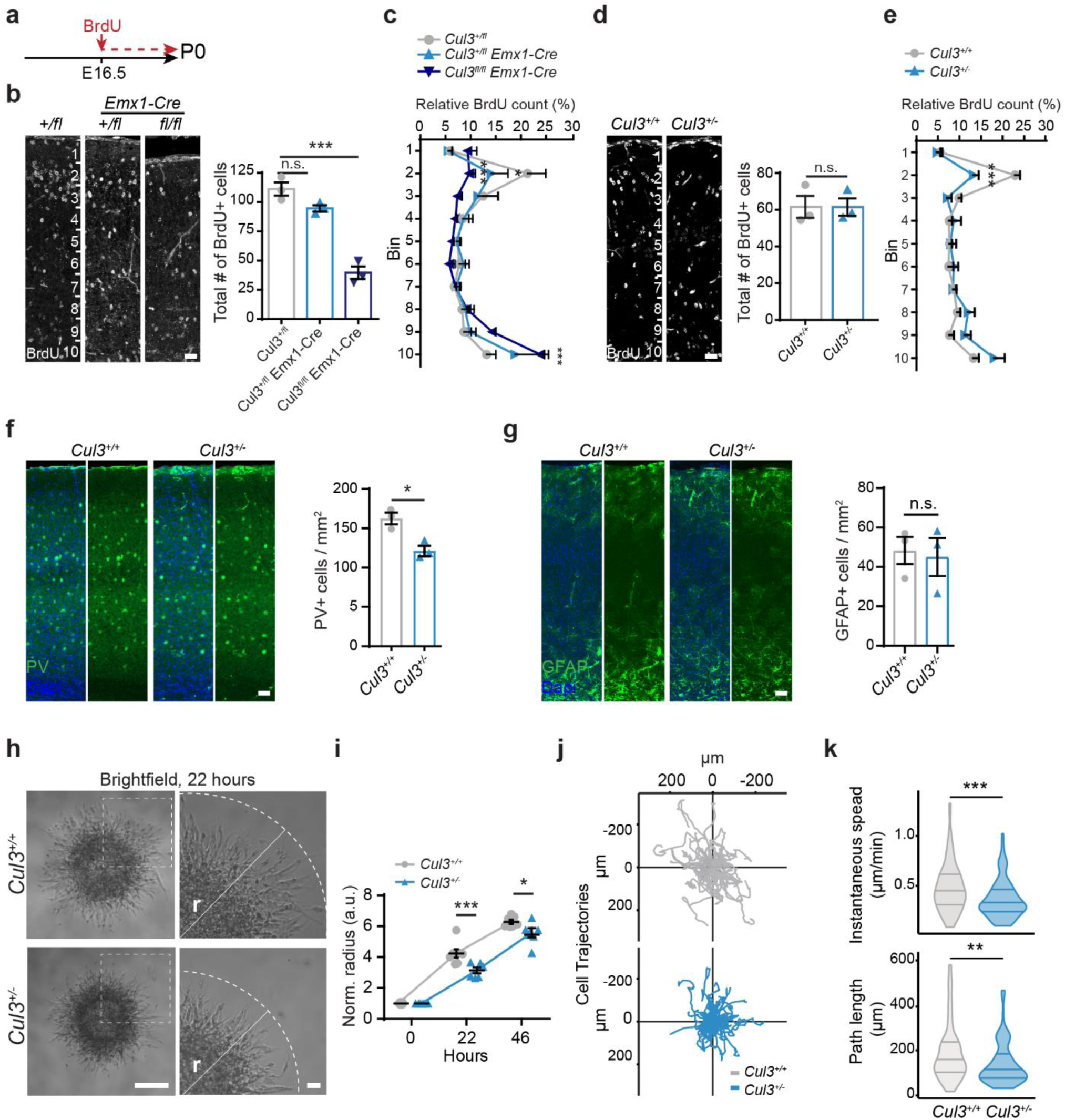
*Cul3* loss leads to migration deficits *in vivo* and *in vitro*. **a**, Scheme of the BrdU birthdate labeling experiments. **b**,**d**, Injection of BrdU at E16.5 and anti-BrdU immunofluorescent stainings and analysis of total BrdU positive (BrdU+) cells in cortical columns at P0 show severe decreased number of BrdU+ cells in *Cul3*^*fl/fl*^ *Emx1-Cre* brains, but not in the *Cul3*^*+/fl*^ *Emx1-Cre* and *Cul3*^*+/-*^ cortex (*n*= 3 littermate pairs pups per genotype; ***P<0.001, n.s. not significant; 1-way ANOVA and Sidak’s multiple comparison test and unpaired two-tailed t-test). **c**,**e**, Bin-wise analysis of relative numbers of BrdU+ cells showed decreased numbers of BrdU+ cells in upper bins and increased numbers of BrdU+ cells in lower bins of all mutant genotypes, *Cul3*^*+/fl*^ *Emx1-Cre, Cul3*^*fl/fl*^ *Emx1-Cre* (c) and *Cul3*^*+/-*^ (e) (*n*= 3 littermate pairs per genotype; *P<0.05, ***P<0.001; 2-way ANOVA and Sidak’s multiple comparison test). **f**-**g**, Representative images and quantification of immunofluorescent stainings against PV+ interneurons and GFAP+ astrocytes in adult coronal cortical sections, showing reduced numbers of PV+ cells (f) but not astrocytes (g) in mutant animals (*n*= 3 littermate pairs per genotype; *P<0.05, n.s., not significant; unpaired two-tailed t-tests). **h**-**i**, *In vitro* migration assay of matrigel embedded neurospheres generated from *Cul3*^*+/+*^ and *Cul3*^*+/-*^ NPCs reveals decreased migratory abilities (r= radius of furthest migrated cell) 22 hours (i, representative images) and 46 hours after plating. Radius was normalized to initial sphere size; (*n*(spheres)= 7/6 *Cul3*^*+/+*^ and *Cul3*^*+/-*^ respectively; *P<0.05, ***P<0.001; 2-way ANOVA and Sidak’s multiple comparison test); **j**-**k**, Cell tracks of *Cul3*^*+/+*^ and *Cul3*^*+/-*^ NPCs detaching from the neurosphere into embedding bovine collagen matrix (*n*(spheres)= 3 per genotype, *n*(cells)= 30 per replicate) imaged in a single plane. Cell trajectories of each cell fixed at origin plotted in Euclidean plane (j). Mean instantaneous speed (k, top) and total cell path length (k, bottom) quantification (**P<0.01; ***P<0.001; Wilcoxon rank sum test). Data presented as mean ± SEM and scatter dot plot in (b,d,f,g right and i), connected mean ± SEM in (c,e) or violin plots with median and first and third quartiles (k). Scale bars: 25 μm in (b,d), 50 μm in (f,g), 200 μm in (h, overview) and 40 μm in (h, close-up). Detailed statistics are provided in Supplementary Table 1.

**Figure 4.**
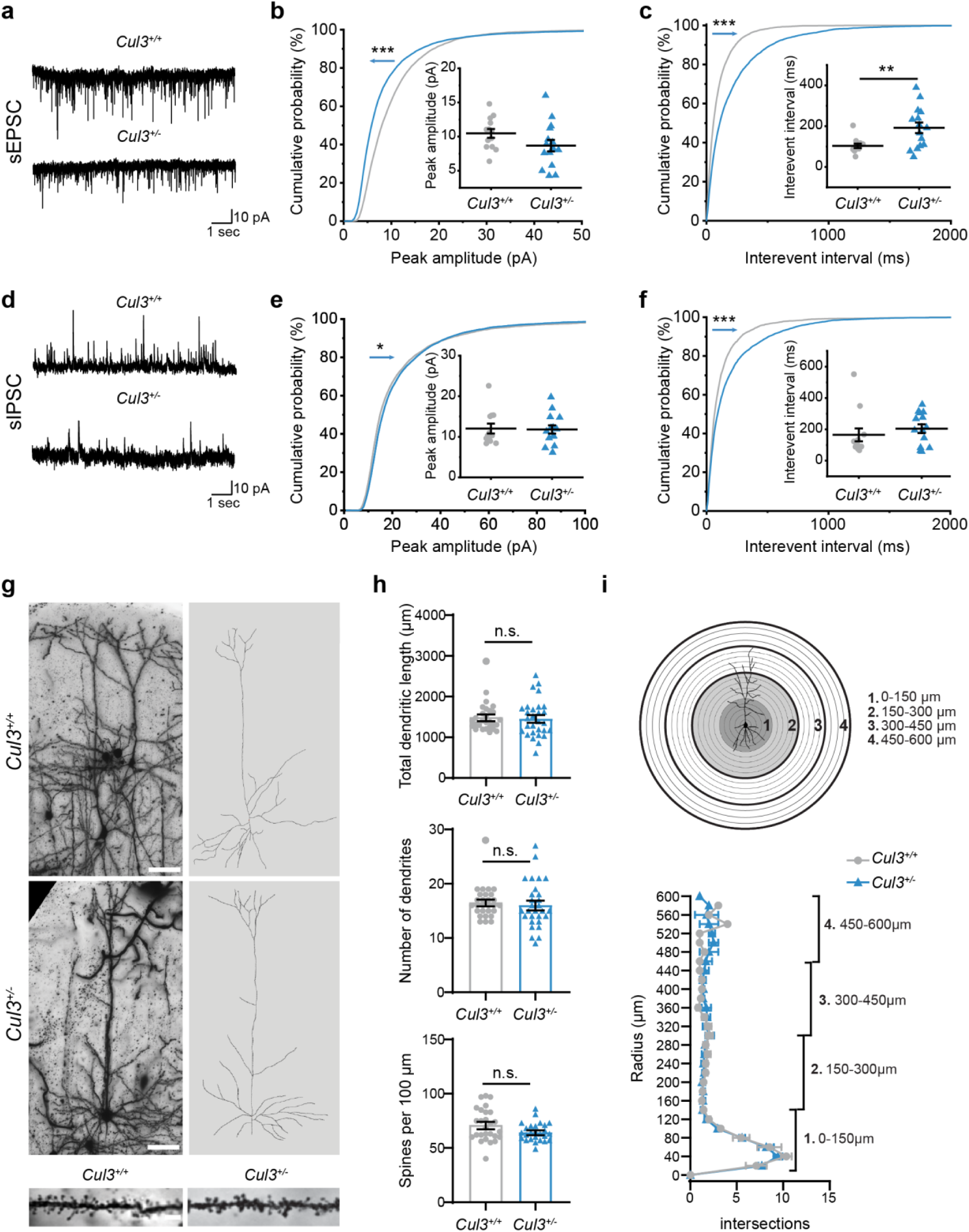
Reduced circuit activity in layer 2/3 pyramidal neurons of adult *Cul3*^*+/-*^ mice, but normal neuronal morphology. **a**, Representative sEPSC traces, recorded by whole cell patch-clamp, at holding potential −70 mV, from layer 2/3 neurons in somatosensory cortex. **b**, Cumulative probability distribution and quantification of sEPSC amplitudes (***p<0.001; Kolmogorov-Smirnov test). **c**, Cumulative probability distribution and quantification of sEPSC interevent intervals (IEI) (*n*(*Cul3*^*+/+*^)=12 cells, *n*(*Cul3*^*+/-*^)= 15 cells from 3 mice respectively; ***p<0.001, **p<0.01; unpaired two-tailed t-test, Kolmogorov-Smirnov test). **d**, Representative sIPSC traces recorded at holding potential +10 mV from layer 2/3 neurons in somatosensory cortex. **e**, Cumulative probability distribution and quantification of sIPSC amplitudes (*p<0.05; Kolmogorov-Smirnov test). **f**, Cumulative probability distribution and quantification of mean sIPSC IEI; (*n*(*Cul3*^*+/+*^)= 12 cells, *n*(*Cul3*^*+/-*^)= 14 cells from 3 mice respectively; ***p<0.001; Kolmogorov-Smirnov test). **g**-**i**, Golgi staining and analysis of the morphology and spine density of layer 2/3 pyramidal neurons in the somatosensory cortex. Brightfield images (g, left) and Imaris reconstructions (g, right) of *Cul3*^*+/+*^ and *Cul3*^*+/-*^ neurons, as well as close-ups of dendrites with spines (g, bottom). Quantification did not reveal any differences in total dendritic length (h, top), number of dendrites (h, center) or spine density (h, bottom) (*n*(cells)= 27-29 from 3 mice per genotype); Sholl analysis was comparable between the mutant and wild type (i, top-scheme; i, bottom-quantification); (*n*(mice)= 3 mice per genotype, at least 9 cells per animal; n.s. not significant; unpaired two-tailed t-tests or two-tailed Mann-Whitney U-test). All analysis was done in adult littermate male mice. Data is shown as mean ± SEM and scatter plots in (b,c,e,f) and connected mean ± SEM in (i). Scale bars: 50 µm in (g, top) and 5 µm in (g, bottom). Detailed statistics are provided in Supplementary Table 1.

We reasoned that complete loss of *Cul3* could exacerbate this phenotype and give us indications about additional *Cul3* deficiency-associated defects. Since constitutive homozygous deletion of *Cul3* is embryonically lethal, we crossed conditional *Cul3* animals (*Cul3*^*fl*^) with an *Emx1-Cre* expressing line, generating forebrain specific heterozygous and homozygous deletions of *Cul3 (Cul3*^*+/fl*^ *Emx1-Cre and Cul3*^*fl/fl*^ *Emx1-Cre* respectively) (Supplementary Fig. 5a). Importantly, *Emx1*-driven *Cre* expression^28^ starts at E10.5, thus inducing *Cul3* deletion at the beginning of forebrain development. While *Cul3*^*+/fl*^ *Emx1-Cre* mice are viable and fertile, like *Cul3*^*+/-*^ animals, *Cul3*^*fl/fl*^ *Emx1-Cre* pups are much smaller than controls (Supplementary Fig. 5b) and die before weaning. In addition, *Cul3*^*fl/fl*^ *Emx1-Cre* mice show severe brain malformations with pronounced cortical and hippocampal atrophy (Fig. 2f, Supplementary Fig. 5c). Similar to what we observed in *Cul3*^*+/-*^ animals, fluorescence imaging of Cux1 and Ctip2 distribution revealed lamination defects in *Cul3*^*fl/fl*^ *Emx1-Cre* and *Cul3*^*+/fl*^ *Emx1-Cre* mice (Fig. 2g-i). Thus, proper *Cul3* dosage is essential to guarantee correct brain development in mouse.

**Figure 5.**
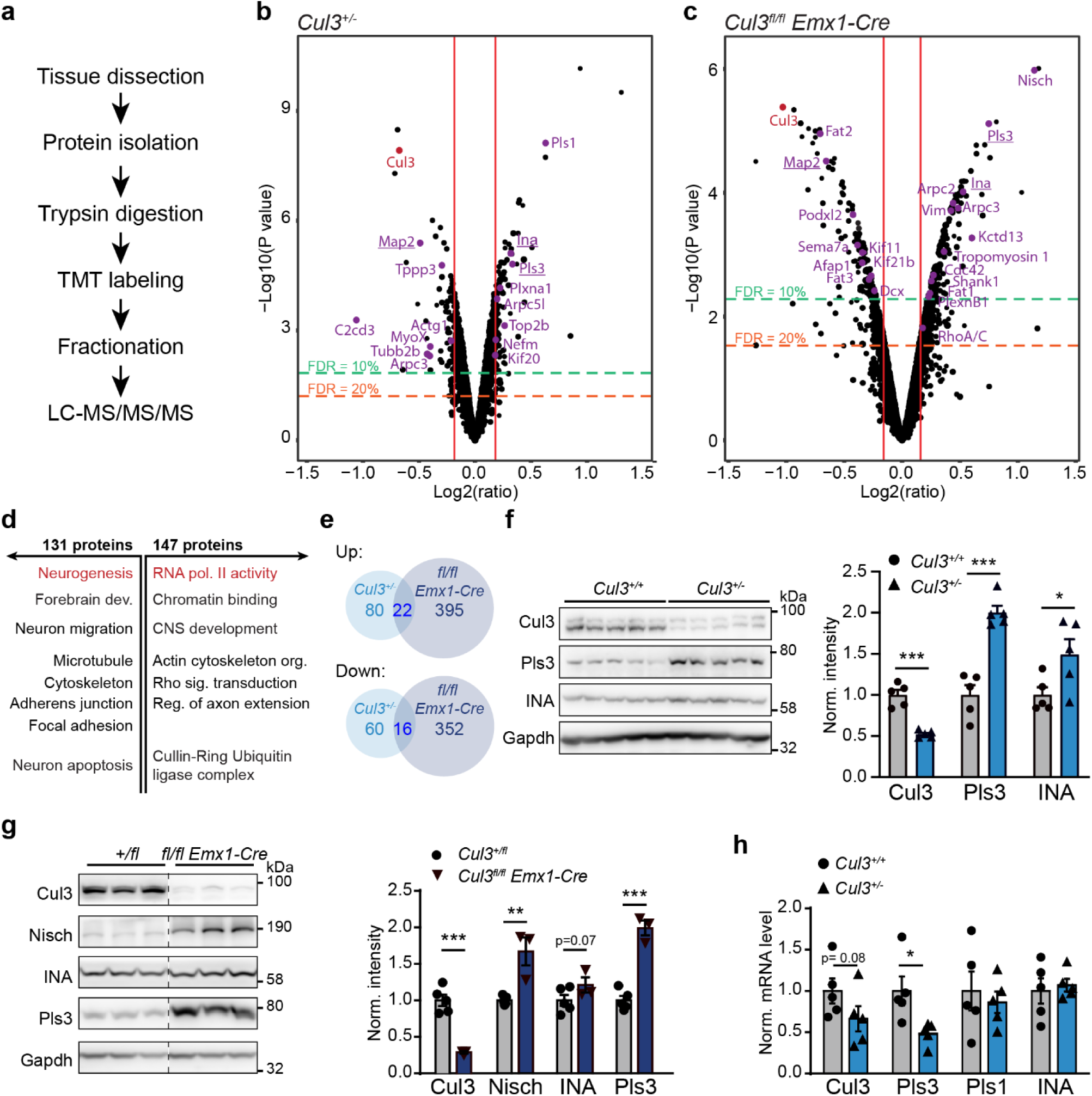
Deregulated cytoskeletal proteins in *Cul3* mutant embryonic forebrain tissue identified by proteomic analysis. **a**, Workflow of sample preparation for proteomic analysis (*n*(*Cul3*^*+/-*^)= 5 per genotype and *n*(*Cul3*^*fl/fl*^ *Emx1-Cre)=* 3 per genotype, male littermate pairs). **b**, Volcano plot of deregulated proteins at 10 and 20% FDR cut-off in the *Cul3*^*+/-*^ developing cortex with 94 (10%) and 102 (20%) up-and 70 (10%) and 76 (20%) down-regulated protein groups. **c**, Volcano plot of deregulated proteins in the *Cul3*^*fl/fl*^ *Emx1-Cre* developing forebrain with 147 (10%), 417 (20%) up- and 131 (10%), 368 (20%) down-regulated protein groups; cytoskeleton related proteins are labeled in purple, Cul3 in red (b,c) (details in Supplementary Tables 3,4). **d**, DAVID functional annotation identified up- (right) and down- (left) regulated proteins of the *Cul3*^*fl/fl*^ *Emx1-Cre* forebrain to be involved in regulating activity of RNA polymerase II, neurogenesis and actin and microtubule cytoskeletal organization (selected GO-terms, significant GO-terms in red: RNA polymerase II core complex, GO: 0005665, adj. p-value= 2.3e-09; Neurogenesis GO:0022008, adj. p-value= 2.7e-05; details in Supplementary Tables 5,6). **e**, Overlap of up- and down-regulated protein groups in the *Cul3*^*+/-*^ and *Cul3*^*fl/fl*^ *Emx1-Cre* cortex at 20% FDR (details in Supplementary Table 7). **f**, Western blot (left) and analysis of Gapdh normalized intensities (right) confirm increased levels of the cytoskeletal proteins Pls3 and INA in *Cul3*^*+/-*^ lysates (*n*= 5 per genotype; *P<0.05, ***P<0.001; unpaired two-tailed t-tests). **g**, Western blot and analysis of Gapdh normalized intensities confirm increased levels of the cytoskeletal proteins Nisch, Pls3 and INA in *Cul3*^*fl/fl*^ *Emx1-Cre* cortical lysates (samples on the same membrane cut for better visualization, *n*(*Cul3*^*+/fl*^)= 5, *n*(*Cul3*^*fl/fl*^ *Emx1-Cre)=* 3; **P<0.01, ***P<0.001; unpaired two-tailed t-tests). **h**, Quantitative real-time PCR analysis of mRNA levels of *Pls1, Pls3* and *INA* normalized to wild-type levels (ΔCq expression values; *n*= 5 per genotype; *P<0.05; unpaired two-tailed t-tests). Data presented as mean ± SEM in (f-h). Detailed statistics are provided in Supplementary Table 1.

### Loss of *Cul3* leads to neuronal migration defects in mice

To identify the origin of the anatomical abnormalities observed in *Cul3*^*+/-*^, *Cul3*^*+/fl*^ *Emx1-Cre* and *Cul3*^*fl/fl*^ *Emx1-Cre* brains we studied cell proliferation, apoptosis, and migration in the three different genotypes, focusing on the time window with highest *Cul3* expression (i.e. E14.5-16.5). First, we injected pregnant females with BrdU at E16.5 and collected the brains of the *Cul3*^*+/fl*^ *Emx1-Cre, Cul3*^*fl/fl*^ *Emx1-Cre* and control embryos two hours after injection. Brain samples were sliced and stained for Sox2, a neural stem cell marker; cleaved caspase-3 (cl. Casp3), a marker for apoptotic cells; and BrdU, as indicator of cells in S-phase (Supplementary Fig. 5d-g). While in *Cul3*^*fl/fl*^ *Emx1-Cre* we found a substantial increase of apoptotic cells in the developing cortex (of which 69.75% ± 1.47 were localized in the ventricular zone) and a corresponding reduction in Sox2+ and BrdU+ cells, we did not observe these anomalies in *Cul3*^*+/fl*^ *Emx1-Cre* animals. This is in line with the more severe phenotype observed in *Cul3*^*fl/fl*^ *Emx1-Cre* mice and with the cell cycle regulation defects described in *Cul3*^*-/-*^ animals^17^.

To test whether cell migration is affected in heterozygous and homozygous *Cul3* mutant mice we again pulsed E16.5 embryos with BrdU, but this time analyzed the number and position of BrdU+ cells in the cerebral cortex at P0 (Fig. 3a). We found a severe reduction in BrdU+ cells in *Cul3*^*fl/fl*^ *Emx1-Cre* P0 animals compared to control samples from *Cul3*^*+/fl*^ littermates (Fig. 3b), consistent with the abovementioned increase in neural cell apoptosis in *Cul3*^*fl/fl*^ *Emx1-Cre* embryos. In addition, we found that a substantially smaller fraction of BrdU+ cells reaches the upper cortical layers and that a significant number of BrdU+ cells remain stranded in lower cortical layers in *Cul3*^*fl/fl*^ *Emx1-Cre* animals (Fig. 3c). These results suggest that complete deletion of *Cul3* in the forebrain leads to neural cell apoptosis and neuronal migration defects. Importantly, while *Cul3*^*+/fl*^ *Emx1-Cre* samples do not show a reduction in total BrdU+ cells, suggesting normal production and survival of cortical neural cells, *Cul3*^*+/fl*^ *Emx1-Cre* pups present a clear reduction of BrdU+ cells reaching the upper part of the cortex (Fig. 3c). We observed a similar defect in the cerebral cortex of constitutive *Cul3*^*+/-*^ mice (Fig. 3d,e), indicating that *Cul3* haploinsufficiency is associated with a neuronal migration phenotype, thus explaining the observed lamination defects.

Next, since we had predominantly quantified the position of glutamatergic neurons, we tested whether other cell types in the brain might be similarly affected. To this end, we counted the density of interneurons, astrocytes and microglia in the adult somatosensory cortex. Interestingly, while the number of interneurons in the cerebral cortex is significantly reduced (Fig. 3f), the amounts of astrocytes and microglia are unchanged in *Cul3* mutant animals (Fig. 3g, Supplementary Fig. 5h,i). This difference may be explained by the fact that *Cul3* expression is highest in excitatory and inhibitory neurons, potentially making them more susceptible to defective protein homeostasis (Supplementary Fig. 3d,g).

To better understand how *Cul3* mutations affect cell migration, we switched to an *in vitro* model system. Analysis of neural progenitor cells (NPCs) generated from E13.5 *Cul3*^*+/+*^ and *Cul3*^*+/-*^ cortices (Supplementary Fig. 5j) confirmed abnormal cell motility *in vitro* (Fig 3h-k, Supplementary Fig. 5k). Specifically, we traced the movement of NPCs moving away from neurospheres over the course of several hours and found that *Cul3* mutant cells travel shorter distances than control cells (Fig. 3h-i). In addition, live imaging analysis revealed that mutant NPCs do not move far from the sphere, migrate less and have reduced migration speed, thus indicating that *Cul3* haploinsufficiency is associated with cell intrinsic defects of migration (Fig. 3j,k, Supplementary Fig. 5k, Supplementary video 1,2).

### *Cul3* haploinsufficiency leads to abnormal neuronal network activity

We reasoned that defects in neural cell migration and cortical lamination could have an important impact on neuronal network activity. Indeed, other ASD-risk genes associated with defects in neuronal migration have been shown to substantially modify neuronal network activity *in vivo*^29^. Spontaneous postsynaptic currents (sPSC) depend both on spiking activity of neurons and spontaneous release of presynaptic vesicles and they are useful for evaluating the global connectivity in the network. To this end, we evaluated the spontaneous synaptic activity in two-months old *Cul3*^*+/-*^ mice, recording from pyramidal neurons in layer 2/3 of the somatosensory cortex in whole-cell configuration. Both spontaneous excitatory and inhibitory post synaptic currents (sEPSC and sIPSC, respectively) are reduced in mutant animals (Fig. 4a-f). In particular, *Cul3*^*+/-*^ mice show a strong reduction in sEPSC frequency, evidenced by an increased mean interevent interval (IEI) (p= 0.005, t-test; cumulative distribution: p<0.0001, Kolmogorov-Smirnov test; Fig.4c) compared to wild-type littermates. The effect of *Cul3* haploinsufficiency on glutamatergic current amplitudes is not evident from the average peak currents (p= 0.104, t-test), however, the cumulative distribution showes a shift of sEPSC toward smaller amplitudes (p<0.0001, Kolmogorov-Smirnov test; Fig. 4b). On the other hand, GABAergic transmission is less affected. *Cul3*^*+/-*^ and wild-type mice showed similar sIPSC peak currents (mean peak: p= 0.89, t-test) even if we notice a slight shift of the cumulative distribution towards higher amplitudes (cumulative distribution: p= 0.02, Kolmogorov-Smirnov test; Fig. 4e) and mean frequency (mean IEI: p= 0.43, t-test; Fig. 4f). Of note, sIPSC distribution is slightly shifted towards lower frequency (IEI cumulative distribution: p<0.0001, Kolmogorov-Smirnov test; Fig. 4f). To test whether the observed differences in neuronal network activity are due to morphological defects, we performed Golgi stainings and analyzed the morphology of layer 2/3 pyramidal neurons in adult *Cul3*^*+/-*^ mice (Fig. 4g-i). However, neither the dendritic length, nor the number of dendrites or spines (Fig. 4h), nor dendritic branching (Fig. 4i) are altered in *Cul3* haploinsufficient mice. Altogether these results point to a tissue level reduction in network activity and global synaptic transmission, likely linked to the cortical lamination defects in *Cul3*^*+/-*^ mice.

### Whole proteome analysis reveals abnormal amounts of cytoskeletal proteins in *Cul3* mutant mice

To gain insight into the molecular mechanisms underlying the observed defects, and in view of Cul3’s E3 ubiquitin ligase function, we assessed the impact of *Cul3* loss on the global proteome of the developing forebrain in *Cul3*^*+/-*^ and *Cul3*^*fl/fl*^ *Emx1-Cre* mutants. Protein extracts from dissected E16.5 cortices from control and mutant animals were analyzed by quantitative proteomics (Fig. 5a). Analysis of the total proteome of *Cul3*^*+/-*^ and *Cul3*^*fl/fl*^ *Emx1-Cre* mutants, as well as corresponding controls, resulted in the identification of 6067 protein groups. For differential protein expression analysis, protein groups were filtered based on fold change and False Discovery Rate (FDR) thresholds. Employing two FDR thresholds of 10% and 20% we identified 94 (10%) and 102 (20%) up- and 70 (10%) and 76 (20%) down-regulated protein groups in the *Cul3*^*+/-*^ embryonic cortex (Fig.5b and Supplementary Table 3). As expected from the more severe phenotype of *Cul3*^*fl/fl*^ *Emx1-Cre* mutant pups, a much larger number of deregulated proteins were identified in conditional homozygous knockout embryos (147 (10%), 417 (20%) up- and 131 (10%), 368 (20%) down-regulated protein groups, Fig. 5c and Supplementary Table 4). Overall, and in agreement with previous observations^30-32^, the observed fold changes were mild, in line with the hypothesis that the ubiquitylated isoform of a protein represents only a small fraction of the total pool of that given protein at any time point^33^. We noticed, however, that proteins with abnormal levels in *Cul3* homozygous mutants are significantly enriched for high confidence ASD-risk genes (*Cul3*^*fl/fl*^ *Emx1-Cre* at 20% FDR, 65 genes, p= 0.0329). The small number of deregulated proteins in the forebrain tissue of *Cul3* haploinsufficient embryos did not justify GO-term enrichment analysis. Therefore, to get an indication of the classes of proteins affected in *Cul3* mutants we performed GO-term enrichment analysis on the *Cul3*^*fl/fl*^ *Emx1-Cre* data only. We found that deregulated proteins were significantly enriched for DNA-directed RNA polymerase II core complex members and proteins involved in neurogenesis. In addition, deregulated proteins were functionally linked to chromatin binding, neuronal migration, apoptosis, actin- and microtubule-cytoskeleton organization and cell adhesion (Fig. 5d, Supplementary Table 5,6). Given the migration phenotype observed in both heterozygous and conditional homozygous mutant animals, we decided to investigate changes in cytoskeletal proteins further and found several of these to be misregulated also in *Cul3*^*+/-*^ embryonic tissue (Fig. 5b,c purple). We annotated these proteins manually, drawing on published literature and focusing on those appearing to follow a dose dependent response to *Cul3* loss (control - *Cul3*^*+/-*^ - *Cul3*^*fl/fl*^ *Emx1-Cre*) (Fig. 5e, Supplementary Table 7). Several have a putative or a confirmed function in cytoskeletal organization, and/or cell migration regulation and differentiation. Of those, some down-regulated proteins are prominent members of a family of microtubule-associated proteins linked to abnormal brain development and cortical lamination defects in humans (i.e. Tubb2b, Dcx)^34,35^. However, as Cul3 targets proteins for degradation, we were most interested in the identified up-regulated proteins, as their up-regulation likely represents a direct consequence of *Cul3* loss. To validate the results obtained by proteomic analysis we selected two of the top up-regulated cytoskeletal-associated proteins observed in both *Cul3*^*+/-*^ and *Cul3*^*fl/fl*^ *Emx1-Cre* samples, Plastin 3 (Pls3) and Internexin neuronal intermediate filament protein A (INA) (Supplementary Table 3,4), and quantified them by western blot. Indeed, we confirmed that *Cul3* deficiency leads to higher amounts of Pls3 and INA in both *Cul3*^*+/-*^ and *Cul3*^*fl/lf*^ Emx1-Cre E16.5 cortical lysates (Fig. 5f,g). Furthermore, we found that the up-regulation of these proteins is not due to increased gene expression since there were no corresponding increases at the mRNA level (Fig. 5h), but rather to a translational or post-translational effect. Interestingly, *Pls3* was actually down-regulated at the mRNA level, likely a compensatory response to the accumulation of the protein, indicating a feedback loop to adjust the protein level of Pls3 and thus pointing towards a potentially important functional role of Pls3 in the brain.

In contrast to early development but in line with an important regulatory function of *Cul3* mostly during early brain development, we found that in adult mutant cortical, hippocampal, and cerebellar tissues, *Cul3* deficiency results in a much smaller number of deregulated proteins (at 10% FDR, cortex: 46 up- and 49 down-, hippocampus: 19 up- and 28 down-, and cerebellum: 29 up- and 42 down-regulated protein groups; Supplementary Fig. 6 and Supplementary Table 8-10). Nevertheless, in samples obtained from adult animals we again found an association between *Cul3* mutations and deregulation of several cytoskeletal-associated proteins also identified at the embryonic time point (e.g. Vim, Pls3). Interestingly, very few protein groups were consistently deregulated in all tissues, suggesting that *Cul3* deficiency has a pleiotropic effect.

**Figure 6.**
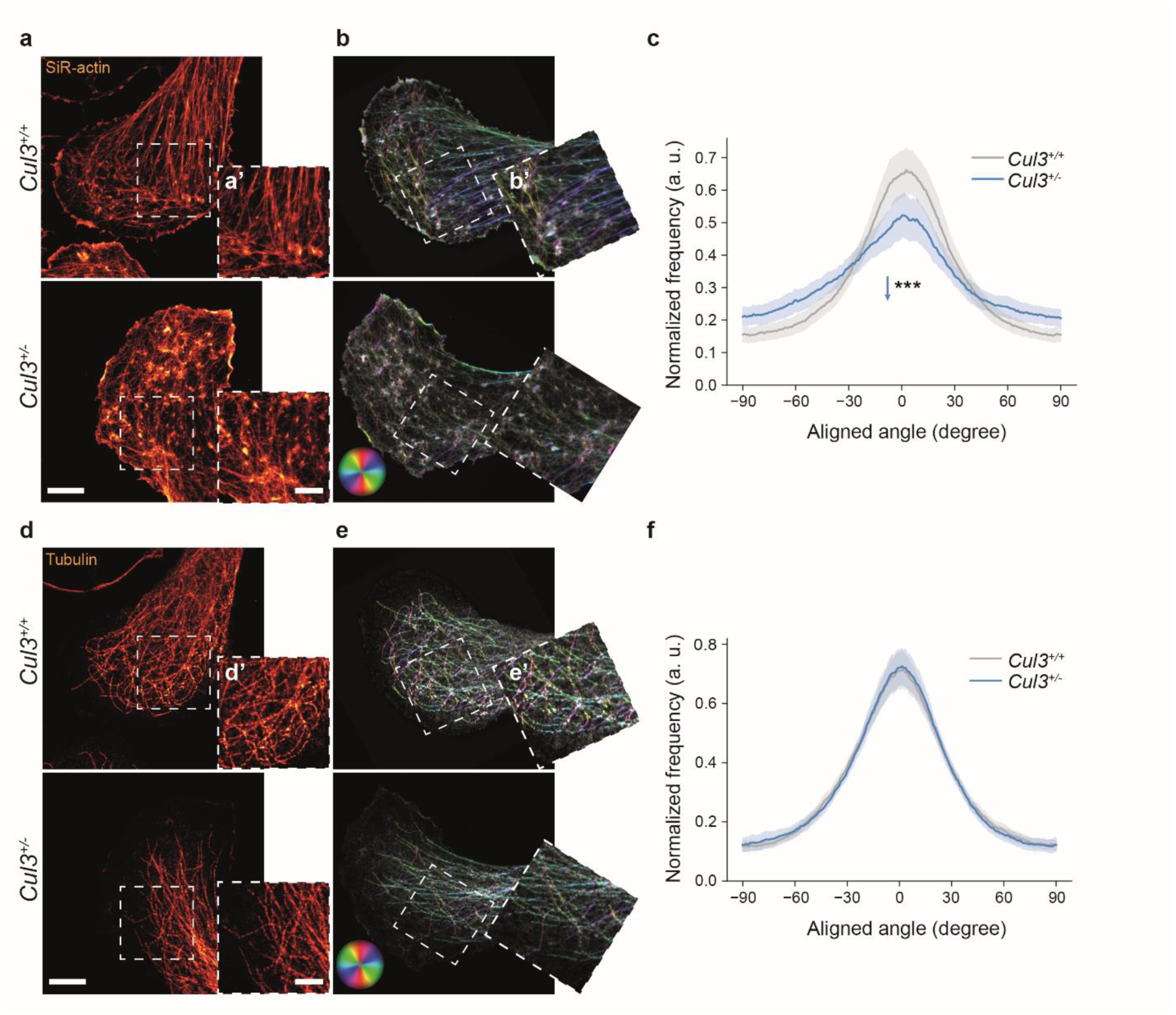
Actin cytoskeleton is disorganized in *Cul3* haploinsufficient NPCs. **a**-**f**, NPCs cultured on Poly-L-ornithine/Laminin coated coverslips were stained using SiR-actin (a) and an anti-tubulin antibody (d). Leading edges of cell protrusions were imaged employing STED-microscopy (close-up images in insets a’,d’). **b**,**e**, Processed, rotated and analyzed images, color code: hue as the orientation angle, saturation as coherency and brightness represents photon counts in the STED image (close-ups in insets b’,e’). **c**,**f**, The dominant orientation is computed as the average orientation angle inside the cell and the mean dominant orientation distribution for actin (c) and microtubules (f) is shown for *Cul3*^*+/+*^ and *Cul3*^*+/-*^ cells (*n*(cells)= 43 per genotype from three independent NPCs preparations; ***P<0.001; two-tailed Welch’s t-test); Plotted is the average angle distribution per group ± 95% confidence intervals. Scale bars: 5 µm in (a,d), 2.5 µm in (insets a’,d’). Detailed statistics are provided in Supplementary Table 1.

However, when we compared our data-set with recently published proteomic analysis of conditional *Cul3* knockouts at adult stages^31,32^, we made a few interesting observations. First, in all data-sets we found down-regulation of Map2, a microtubule associated protein and known component of the neuronal cytoskeleton. Reduced levels of Map2 may be due to the reduced cortical layer thickness and numbers of neurons. Second, we found that the only protein consistently up regulated in mutant animals is Pls3, a still poorly characterized actin bundling protein^36,37^. Pls3 is altered in the developing and adult cortex, the adult hippocampus, but not in the adult striatum^31,32^. Interestingly, a pathogenic variant of *PLS3* has recently also been described in a patient with idiopathic osteoporosis and ASD^38^. Thus, our analysis points to a central role of Cul3 in the homeostatic regulation of cytoskeletal proteins of which the most significant might be Plastin 3.

### Abnormal actin organization in *Cul3* haploinsufficient cells

Proteomic analysis of embryonic cortices highlighted an alteration of cytoskeletal protein levels in *Cul3* mutant samples. In particular, a number of actin binding proteins were significantly more abundant in cortices obtained from *Cul3* mutant animals than in wild types. To assess whether abnormal homeostasis of actin cytoskeleton proteins results in structural abnormalities in moving cells^39^ we analyzed actin conformation in neural stem cells *in vitro* employing stimulated emission depletion (STED) super-resolution imaging to resolve the intricate actin network at the leading edge. Specifically, we stained control and *Cul3* mutant NPCs with SiR-actin, a fluorescent probe for F-actin, as well as a tubulin antibody and analyzed actin filament orientation at the leading edge in diffraction-unlimited images. We found that *Cul3* deficiency leads to a disorganized actin-architecture in the leading edge of adherent *Cul3*^*+/-*^ NPCs (Fig. 6a-c), while the microtubule organization appears normal (Fig. 6d-f). Decreased directionality of actin filaments may in principle be caused by an increased number of focal adhesion sites, as these puncta contain all possible angles and would thereby decrease the measured dominant direction. Thus, we further analyzed the number of adhesion points in control and mutant cells. Surprisingly, adhesion site counting revealed a slight, yet significant, decrease in focal adhesions in the mutant cells (Supplementary Fig. 7a-c). A reduction of adhesion points is also in line with our proteomic data, which identified a reduction of cell adhesion proteins (such as Aatf, Afap1, Bsg, Cd151). Taken together, high resolution imaging indicates a disorganized actin mesh at the leading edge, probably constituting the underlying cell biological correlate of the migration defects observed in *Cul3* mutant cells.

**Figure 7.**
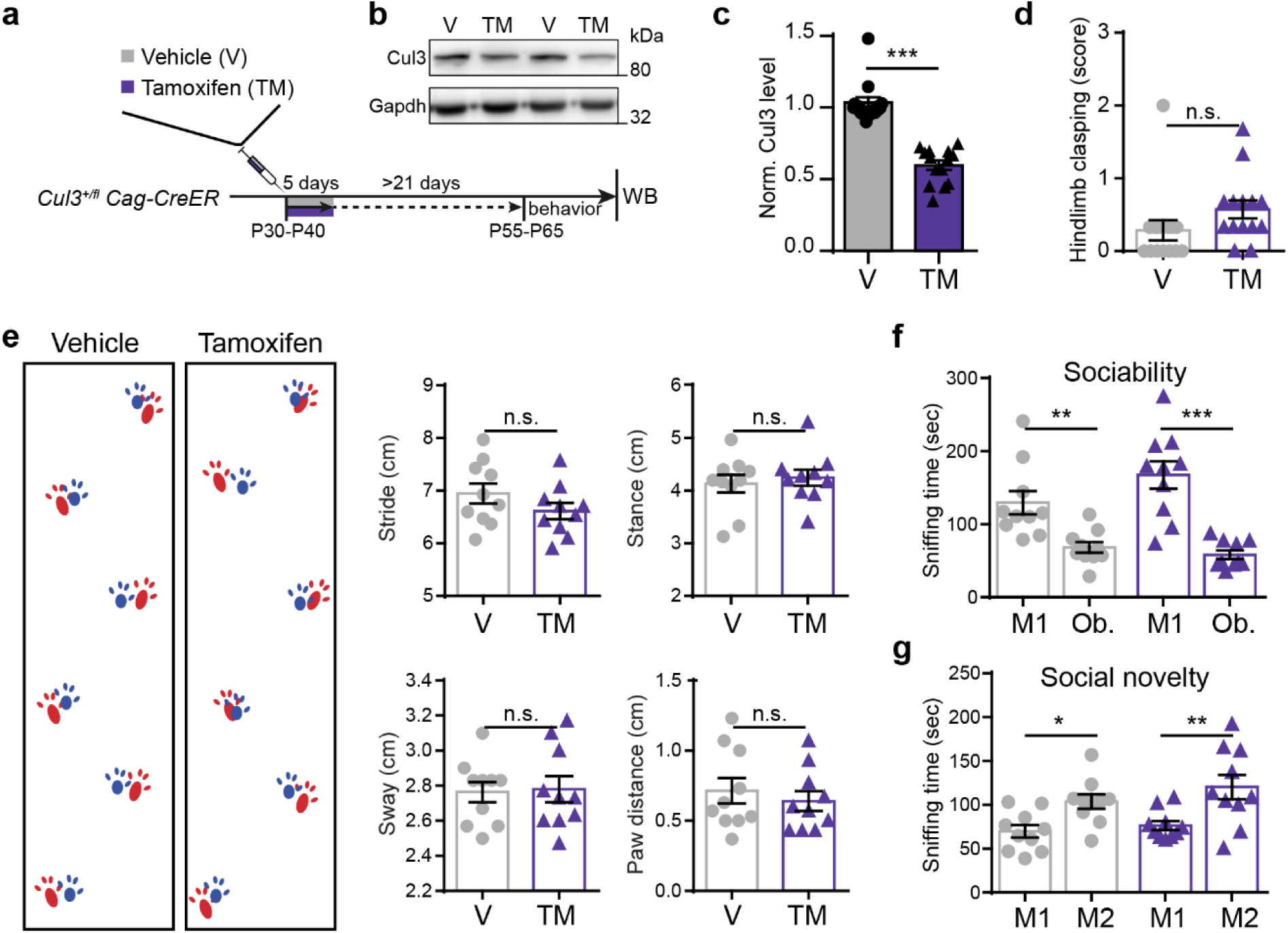
*Cul3* loss after completion of main developmental milestones does not lead to behavioral abnormalities in mice. **a**-**b**, 30 to 40 day-old *Cul3*^*+/fl*^ *Cag-CreER* double transgenic mice were injected for 5 consecutive days with either 100 mg/kg body weight tamoxifen (TM) or vehicle (V, corn oil). Behavioral tests were performed after at least 21 days post-last injection (P55-65), followed by Western blot analysis of Cul3 levels in the brain (a, scheme and b, representative Western blot). **c**, Quantification of normalized Cul3 protein levels in the brain of tamoxifen treated mice, normalized to the levels of vehicle injected controls (*n*= 14 per condition, sex-matched littermate pairs; ***P<0.001; unpaired two-tailed t-test). **d**, Hindlimb clasping scoring from 0-1 (normal) to 3 (most severe) did not reveal any difference between conditions (*n*= 14 per condition, sex-matched littermate pairs; n.s. not significant; unpaired two-tailed t-test). **e**, Gait analysis of tamoxifen treated and vehicle treated mice (representative foot-prints, left), did not reveal any difference between conditions, in stride, stance, sway or paw distance (*n*= 10 mice per condition, sex-matched littermate pairs; n.s., not significant, unpaired two-tailed t-tests). **f**-**g**, *Cul3* loss at P30-P40 does not induce abnormal behavior in the three chamber sociability test, both tamoxifen and vehicle treated *Cul3*^*+/fl*^ *Cag-CreER* mice significantly prefer a stranger mouse (M1) over the caged object (Ob.) (f), and the novel stranger (M2) over the already familiar mouse (M1) (g) (*n*= 10 mice per condition, sex-matched littermate pairs; *P<0.05, **P<0.01 and ***P<0.001; 1-way ANOVA and Sidak’s multiple comparison test). Data is presented as mean ± SEM in scatter dot plots. Detailed statistics are provided in Supplementary Table 1.

### Behavioral abnormalities are associated with *Cul3* developmental functions

The point(s) in time when ASD mutations exert their effects on the brain remains elusive in most cases, as is also their impact on the appearance of ASD core features. However, identifying these critical temporal windows may be essential to properly design therapeutic strategies and clinical trials. In order to understand how critical the loss of *Cul3* is at developmental stages, we investigated the link between the observed embryonic cellular defects and the appearance of mouse behavioral phenotypes by analyzing the effects of *Cul3* deletion at a later time point. To induce deletion of *Cul3* postnatally, we crossed our conditional *Cul3* allele (*Cul3*^*+/fl*^) with animals expressing a tamoxifen-responsive Cre recombinase (*Cag-CreER*). Thus, we induced heterozygous *Cul3* deletion by tamoxifen injections of *Cul3*^*+/fl*^ *Cag-CreER* mice between P30 and P40, and performed behavioral analysis of these animals and vehicle treated littermates at P55-60 (Fig. 7a). Consecutive daily tamoxifen injections (100 mg/kg, five days, starting at P30) significantly decrease *Cul3* protein to about half of control levels in *Cul3*^*+/fl*^ *Cag-CreER* brain tissue (Fig. 7b,c). However, induction of *Cul3* haploinsufficiency at P30 does not result in any obvious behavioral defects. In particular, we did not observe any increase in hindlimb clasping events or abnormalities in gait or social novelty, behaviors which were clearly perturbed in animals with germline *Cul3* haploinsufficiency (Fig. 7d-f). These results indicate that developmental stages are critical for the appearance of *Cul3*-associated behavioral phenotypes and suggest that later interventions may be ineffective in patients carrying mutations in the *CUL3* gene.

## Discussion

*De novo* loF mutations in *CUL3* are an important cause of ASD, motor deficits and intellectual disability in humans. *CUL3* is also involved in the presentation of the 16p11.2 deletion and duplication syndrome, associated with a variety of neurological issues. Therefore, understanding the role of *CUL3* in the mammalian brain is of utmost importance. While the molecular function of CUL3 is well described, its role in the brain, particularly during development, remains largely unclear. To address these questions and model the patients’ *CUL3* haploinsufficiency, we analyzed a *Cul3* construct valid mouse model.

We found that constitutive *Cul3* heterozygous deletion leads to several behavioral abnormalities, including sociability issues, motor dysfunctions, and olfactory hyper-reactivity in adult animals. While this confirms the importance of maintaining correct *Cul3* dosage in the mammalian brain, it also implicates *Cul3* in the function of different brain regions, in line with the complex presentation of *CUL3* mutant patients. Importantly, for our behavioral studies we invested effort in generating and analyzing male and female cohorts separately but we did not observe major phenotypic differences between genotypes of different sexes, supporting the observation that in humans *CUL3* mutations similarly affect males and females. Thus, *CUL3* mutations do not appear to contribute to the skewed sex ratio in ASD cases.

While a gross morphological analysis did not reveal major abnormalities in the adult mutant brain, we found that *Cul3* haploinsufficiency leads to cortical lamination defects. Lamination abnormalities in *Cul3*^+/-^ animals are linked to migration defects, which lead to retention of neuronal cells in lower cortical layers. *Cul3* mutation-associated neuronal migration defects are cell-autonomous and do not depend on brain-specific cues, as we also observe migration phenotypes *in vitro*. Furthermore, our data indicate that *Cul3* deficiency does not affect the number of astrocytes and microglia cells in the cortex, thus suggesting that *Cul3* regulates neuronal-specific processes, possibly due to its much higher expression in neuronal cell types compared to other brain cells types. A role of *Cul3* in cell migration was already hypothesized due to its connection with *Kctd13*, one of the genes localized in the part of the chromosome 16 associated with 16p11.2 deletion syndrome. *Kctd13* encodes a substrate-linking protein for Cul3 and was suggested to bind RhoA, a regulator of the actin cytoskeleton, and thereby targeting it for ubiquitylation and degradation^40^. However, *Kctd13* deletion does not lead to elevated RhoA until after P7 and adult *Kctd13* heterozygous knockout mice do not have major structural brain differences as assessed by MRI^41^. Thus, in agreement with a lack of RhoA increase in *Cul3*^*+/-*^ embryonic forebrain tissue, as seen in our proteomics data, the observed migration and lamination defects are most likely caused by a more general effect of *Cul3* loss on actin cytoskeleton-associated proteins, as indicated by our proteomics analysis at E16.5, rather than driven by RhoA. Integrity of the actin-cytoskeleton is known to be essential in multiple cell types to facilitate cell migration by generating protrusive and contractile forces^42^. Abnormal homeostasis of actin cytoskeletal proteins likely leads to the observed disorganization of actin architecture at the leading edge of migrating cells, thus explaining the migration defects displayed by *Cul3* mutant cells *in vivo* and *in vitro*.

Lamination defects, even when subtle, can have a profound effect on the physiology of the brain and disrupt the stereotyped organization of the microcolumnar structures typical of the neocortex. Accordingly, lamination defects have been previously associated with ASD in human and mouse models due to alteration of neuronal connectivity^43^. Similarly, *Cul3* haploinsufficient mice show decreased spontaneous activity of layer 2/3 cortical excitatory and inhibitory neurons, possibly due to a reduction of total neurons reaching these upper cortical layers and an overall incorrect laminar organization.

Interestingly, complete deletion of *Cul3* leads to additional phenotypes including increased apoptosis during the neural stem cell proliferation phase, possibly due to a defect in cell cycle progression. This is matched by a more severely affected proteostasis in the developing brain of Cul3^*fl/fl*^ *Emx1-Cre* embryos. Defects of cell cycle progression were already associated with complete depletion of *Cul3* and do underlie the lethality of *Cul3* null mice^17^. Thus, studying construct valid *Cul3* models is critical to understanding the bases of ASD in patients, as due to its stronger effect, homozygous deletion can obscure the underlying cellular and molecular drivers of *Cul3* haploinsuffisciency-linked phenotypes.

Importantly, while two recent studies^31,32^ analyzed the effect of *Cul3* deficiency and haploinsufficiency in the adult mouse brain, both studies fell short of studying the developmental consequences of *Cul3* mutations in detail and employed Cre-lines that may miss critical developmental issues. In contrast, here we focused on the constitutive effects of *Cul3* haploinsufficiency and discovered its crucial role in cell migration and proper organization of the cortex. Interestingly, when comparing our proteomic data with these recently published studies we found that the only proteins consistently altered in both these studies and our present investigation are actin and tubulin associated proteins, including Pls3 and Map2. Thus, although there are some discrepancies, probably due to different experimental choices (e.g. mouse model employed, time point analyzed etc.), the combined evidence suggest a role of *Cul3* in regulating cytoskeletal organization. Furthermore, all data sets point to a potential important role of Pls3 in the brain. The function of Pls3 in the brain is mostly unknown but some data suggested that reduced levels of this protein may play a role in motor neuron degeneration in spinal muscular atrophy^44^.

In addition, our observations point to a central role of *Cul3* in early brain development. Conversely, heterozygous deletion of *Cul3* in adult animals does not cause obvious behavioral defects. Therefore, although a direct connection is still missing, it seems plausible that *Cul3* haploinsufficiency-associated migration defects play a central role in the behavioral abnormalities associated with ASD. While some questions regarding the exact temporal trajectory remain to be answered, our findings point to a critical developmental time window for the emergence of *Cul3* related, ASD-linked, behavioral abnormalities. As the field is currently facing major roadblocks in moving promising treatment strategies from preclinical trials to successfully completed clinical trials in humans, stratification of the very heterogeneous patient population into more defined biological subgroups will be necessary. In that light, here we provided novel insights into the pathophysiological and temporal basis of *Cul3-*linked behavioral abnormalities that might one day inform drug development and clinical trial design.

### Accession code

The mass spectrometry proteomics data have been deposited to the ProteomeXchange Consortium via the PRIDE ^45^ partner repository with the data-set identifier PXD017040.

## Acknowledgements

We thank A. Coll Manzano and F. Freeman for technical assistance, S. Deixler, A. Lepold and R. Stemberger for the management of our animal colony, as well as M. Schunn and the Preclinical Facility team for technical assistance. We thank K. Heesom and her team at the University of Bristol Proteomics Facility for the proteomics sample preparation, data generation, and analysis support. Further, we thank M. Sixt for his advice regarding cell migration and the fruitful discussions. This work was supported by the Austrian Science Fund (FWF) to G.N. (W1232-B24) and to J.G.D (I3600-B27).

## Contributions

J.M. designed and performed experiments, analyzed data, and prepared figures. L.A.S., B.B. and S.T. performed experiments and data analysis. A.N., C.S. and C.P.D. analyzed data. C.K., Z.D., L.S.K. and E.C. performed experiments. J.G.D. supervised STED-imaging. G.N. conceived and supervised the study. G.N. wrote the paper together with J.M.. All authors read and approved the final version of the manuscript.

