## Supplemental Figures and Methods for "*Cul3* regulates cytoskeleton protein homeostasis and cell migration during a critical window of brain development"

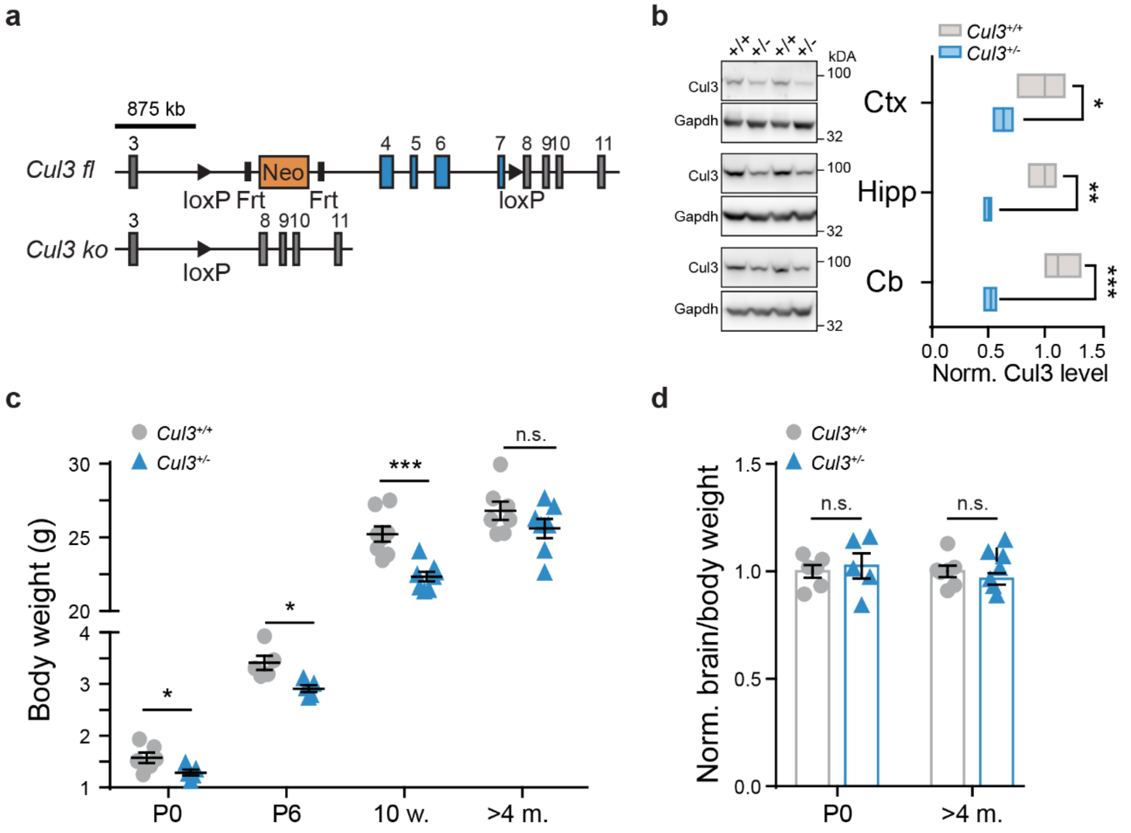


**Supplementary Figure 1. The conditional *Cul3* allele and its deletion in mice. a**, Scheme of the conditional *Cul3* allele in mice in which exons 4-7 are flanked by two loxP sites (*Cul3*$^{fl}$). Cre-mediated recombination leads to genomic excision of the flanked region resulting in the *Cul3* knockout (*Cul3*$^{ko}$) allele. **b**, Representative Western blots and analysis of adult cortex (Ctx), hippocampus (Hipp) and cerebellum (Cb) reveal significantly decreased Cul3 levels in all brain regions of *Cul3*$^{+/-}$ mice (*n*= 3 per genotype; *P<0.05, **P<0.01, ***P<0.001; unpaired two-tailed t-tests). **c**, *Cul3*$^{+/-}$ mice are born with reduced body weight as compared to their wild-type littermates, a growth defect persisting until early adulthood (P6 and 10 weeks), but recovered at 4 month of age (*n*(P0, P6, 10w., 4m.)= 6, 5, 8, 7 per genotype respectively; *P<0.05, ***P<0.001, n.s. not significant; unpaired two-tailed t-tests). **d**, Brain to body weight ratios, normalized to control littermates are normal at P0 and 4 month of age (*n*(P0, 4m.)= 6, 7 per genotype; n.s. not significant; unpaired two-tailed t-tests). Data is presented as mean ± SEM. Detailed statistics can be found in Supplementary Table 1.

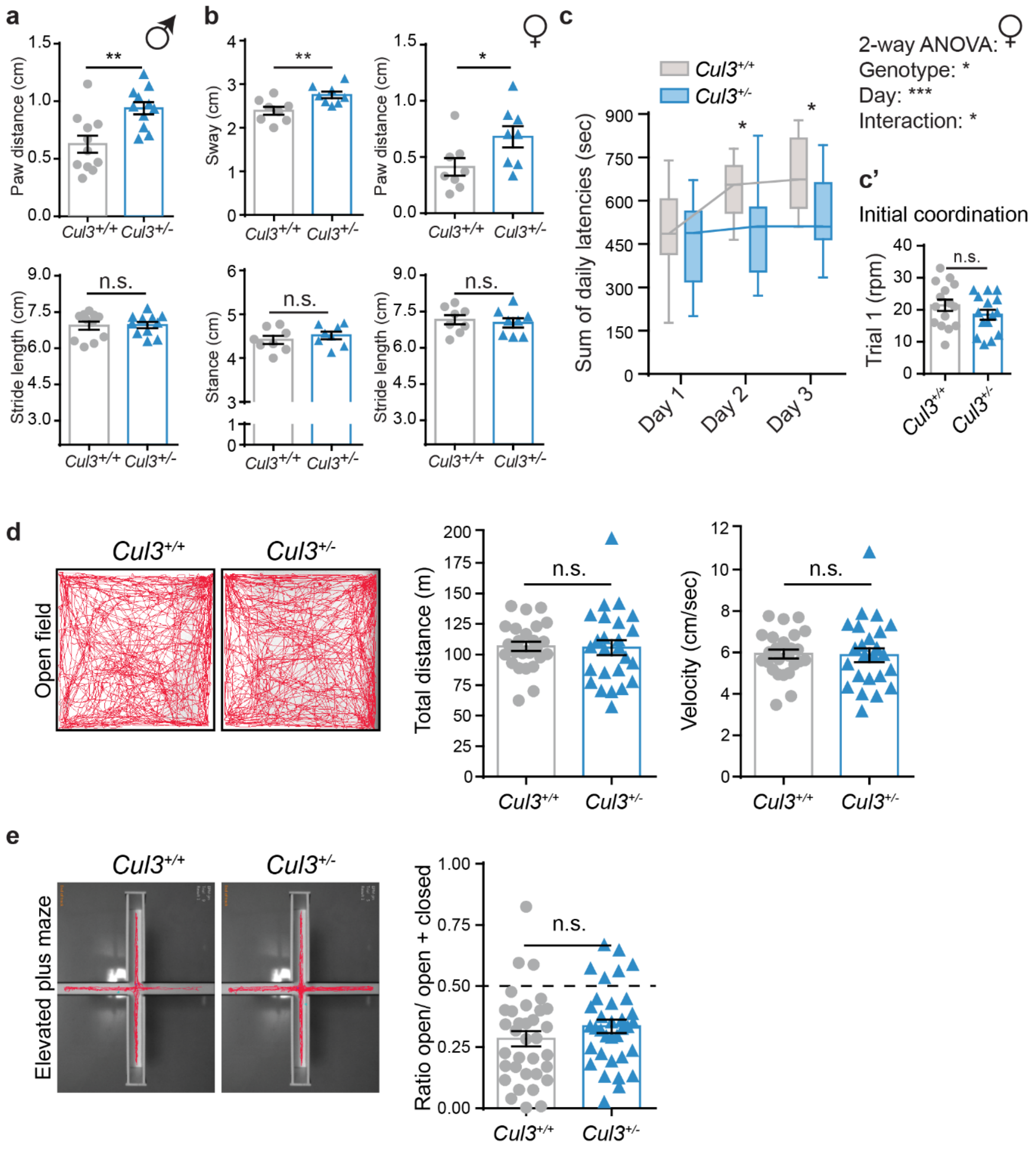


**Supplementary Figure 2. Further behavioral features of *Cul3* haploinsufficient mice. a**, Increased paw distance (top) but normal stride length (bottom) in *Cul3*$^{+/-}$ male animals (*n*= 11 male mice per genotype, littermates; **P<0.01; n.s. not significant; two-tailed Mann-Whitney U test or two-tailed t-test). **b**, Altered gait also in *Cul3* haploinsufficient female mice evidenced by inter-genotype comparison of sway (left top), stance length (left bottom), paw distance (right top) and stride length (right bottom) (*n*= 8 females per genotype, littermates; *P<0.05, **P<0.01, n.s. not significant; unpaired two-tailed t-tests). **c**-**c'**, Accelerating RotaRod test revealing defects in motor learning and coordination also in female *Cul3*$^{+/-}$ mice; shown are the sum of daily latencies of three trials per day on three consecutive days (c) and the final rpm on day one - trial 1, as measure of initial coordination (c') (*n*= 15 female littermate pairs per genotype; *P<0.05, n.s. not significant; 2-way ANOVA and Sidak's multiple comparison test and unpaired two-tailed t-test). **d**, Normal exploratory behavior in the open field. Representative trajectories (left), and quantification of the total distance moved (center) and velocity (right) reveal no differences between *Cul3*$^{+/+}$ and *Cul3*$^{+/-}$ mice (*n*= 25 sex-matched littermate pairs, females (*n*= 13) and males (*n*= 12); n.s. not significant; unpaired two-tailed t-tests). **e**, Performance of *Cul3*$^{+/-}$ mice in the elevated plus maze (total duration 6 min), representative trajectories (left) and quantification of the ratio of time spent on the open/ open+ closed arm (*n*= 34 sex-matched littermate pairs of wild-type and *Cul3*$^{+/-}$ animals, females (*n*= 15) and males (*n*= 19); n.s. not significant; unpaired two-tailed t-test). Data is presented as mean ± SEM. Detailed statistics can be found in Supplementary Table 1.

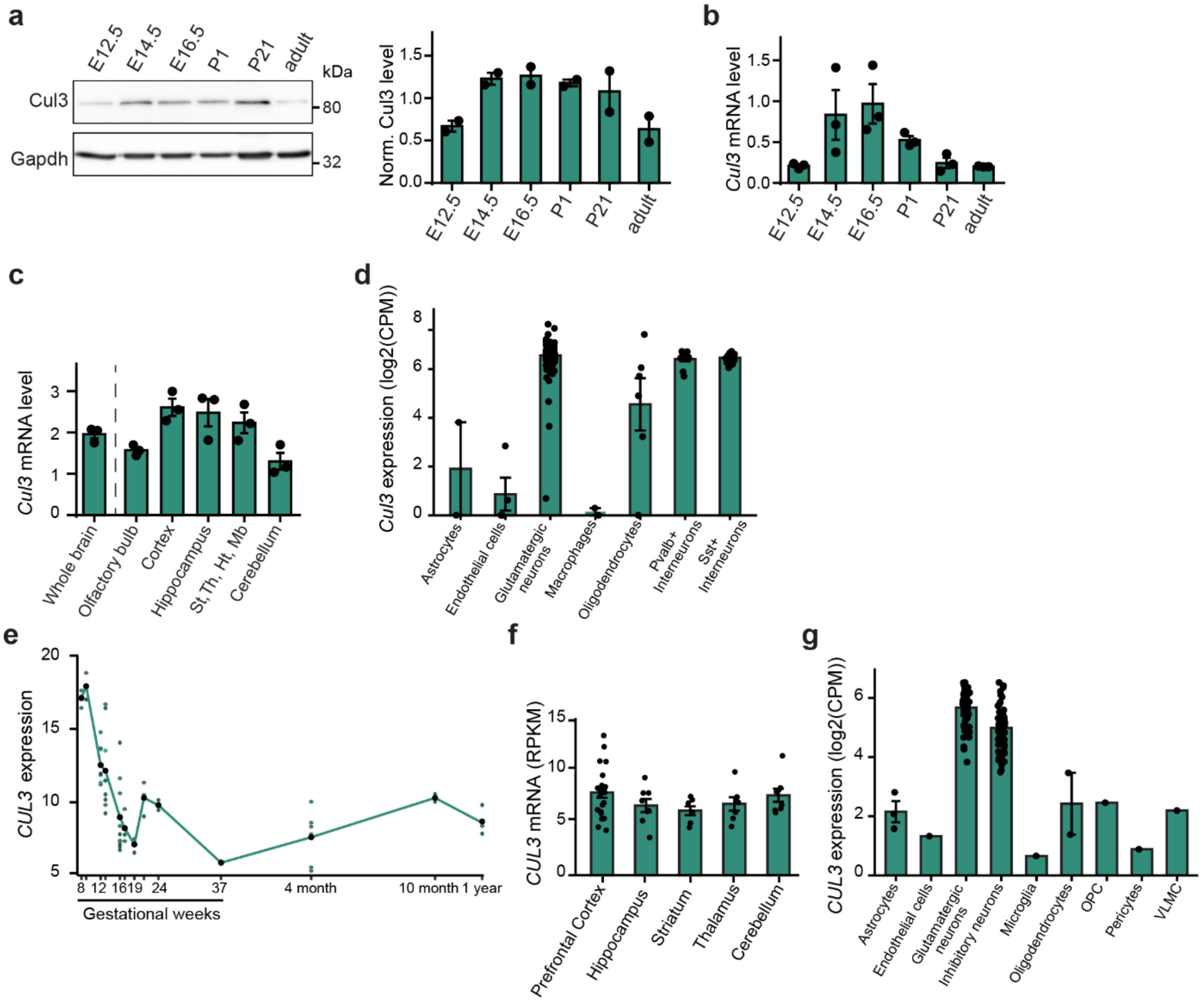


**Supplementary Figure 3. *Cul3* expression peaks during early development in both, mouse and humans. a**, Western blot and quantification of E12.5, E14.5, E16.5, P1, P21 and adult brain lysates of C57BL6J wild-type animals show highest Cul3 protein levels during developmental time-windows important for brain development (*n*(pooled tissue)= 3 animals per time point, *N*(WB)= 2). **b**-**c**, Quantitative real-time PCR analysis of *Cul3*, in brain development (b) and in adult brain regions (c) of C57BL6J wild-type animals, confirms expression peaks during E14.5 and E16.5 and in cortex and hippocampal tissue. Lower *Cul3* levels were observed in the olfactory bulbs and the cerebellum (St= Striatum, Th= Thalamus, Ht= Hypothalamus, Mb= Midbrain) (*n*(tissue)= 3 animals, *N*(qPCR)= 3; ΔCq expression values are plotted). **d**, Normalized *Cul3* expression across cell types in the adult mouse brain based on data from the Allen Cell Types Database [https://portal.brain-map.org/atlases-and-data/rnaseq]. Data points indicate individual cell type clusters that were aggregated for this analysis. **e**, *CUL3* expression in cortical samples across human development based on data from the BrainSpan Atlas [http://www.brainspan.org/static/download.html]. X-axis shows age and y-axis shows expression in RPKM. Data points indicate individual samples. **f**, *CUL3* expression in RPKM across regions of the adult human brain based on data from the BrainSpan Atlas [http://www.brainspan.org/static/download.html]. Data points indicate individual samples. **g**, Normalized *CUL3* expression across cell types in the adult human brain based on data from the Allen Cell Types Database [https://portal.brain-map.org/atlases-and-data/rnaseq]. Data points indicate individual cell type clusters that were aggregated for this analysis. Data is presented as mean ± SEM.

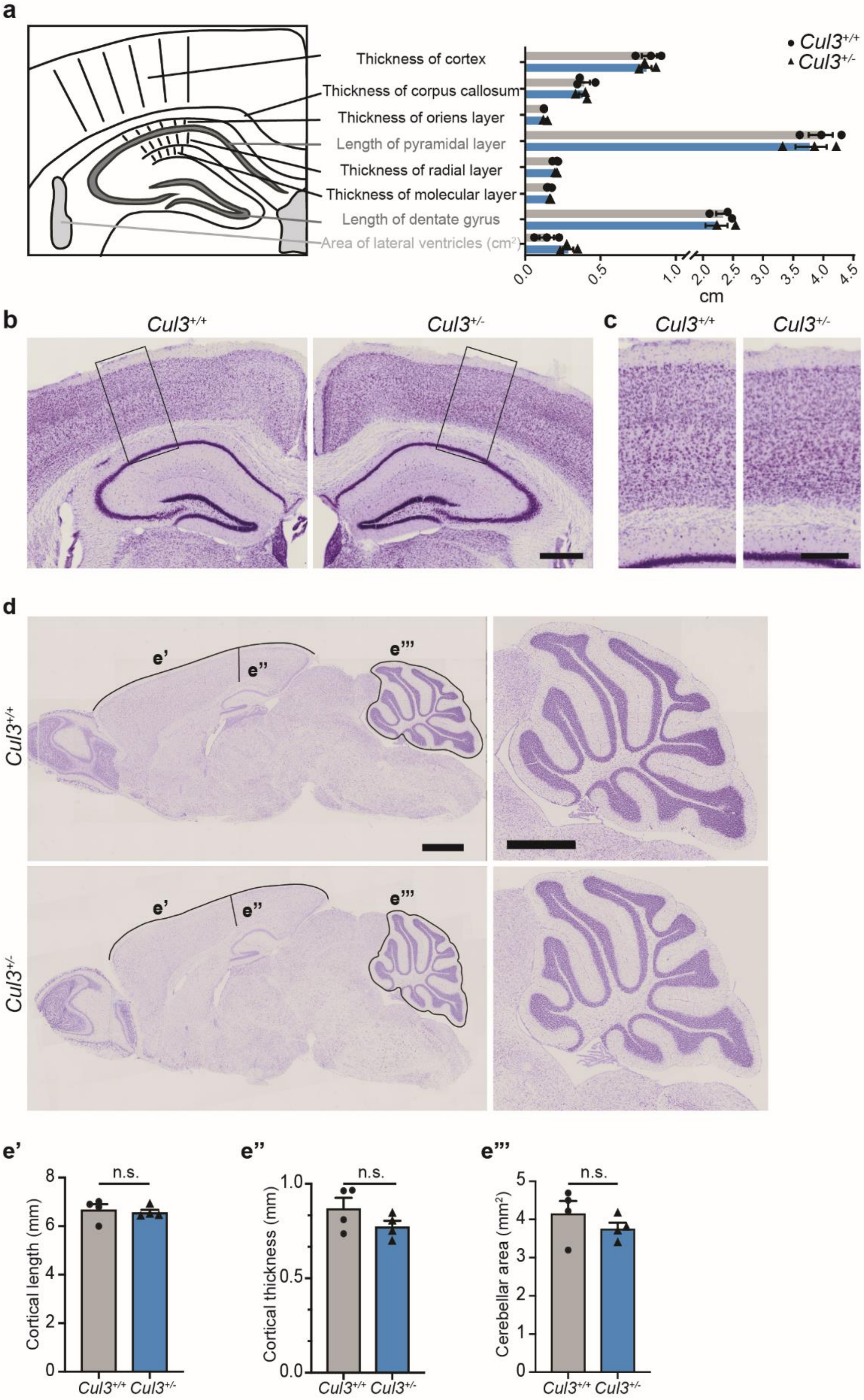
a
Cul3+/+
Cul3+/-
Thickness of cortex
Thickness of corpus callosum
Thickness of oriens layer
Length of pyramidal layer
Thickness of radial layer
Thickness of molecular layer
Length of dentate gyrus
Area of lateral ventricles (cm2)
0.0 0.5 1.0 2.0 2.5 3.0 3.5 4.0 4.5
cm
b
Cul3+/+
Cul3+/-
c
Cul3+/+
Cul3+/-
d
Cul3+/+
Cul3+/-
e'
e''
e'''
e'
Cortical length (mm)
n.s.
Cul3+/+ Cul3+/-
e''
Cortical thickness (mm)
n.s.
Cul3+/+ Cul3+/-
e'''
Cerebellar area (mm2)
n.s.
Cul3+/+ Cul3+/-

**Supplementary Figure 4. Gross brain morphology of *Cul3* haploinsufficient mice appears normal. a**, Scheme of coronal forebrain sections and the measured brain features in adult *Cul3*$^{+/-}$ and *Cul3*$^{+/+}$ littermates revealed no differences between genotypes (*n*= 3 mice per genotype; unpaired two-tailed t-tests). **b**-**c**, Representative Nissl stainings of coronal forebrain sections in mutant and wild-type mice, analyzed in (a) and close-ups of cortical columns in boxed regions in (c). **d**-**e'''**, Overview (left) and cerebellar close-up (right) of Nissl stained adult sagittal brain sections and measurements of cortical length (e'), cortical thickness (e'') and cerebellar area (**e'''**) confirmed these results, despite slight, yet not significant, decreases in the latter could be observed (*n*= 4 mice per genotype, littermates; n.s. not significant; unpaired two-tailed t-tests). Data is presented as mean ± SEM. Scale bars: 500 µm in (b), 250 µm in (c), 1 mm in (d, left and right).

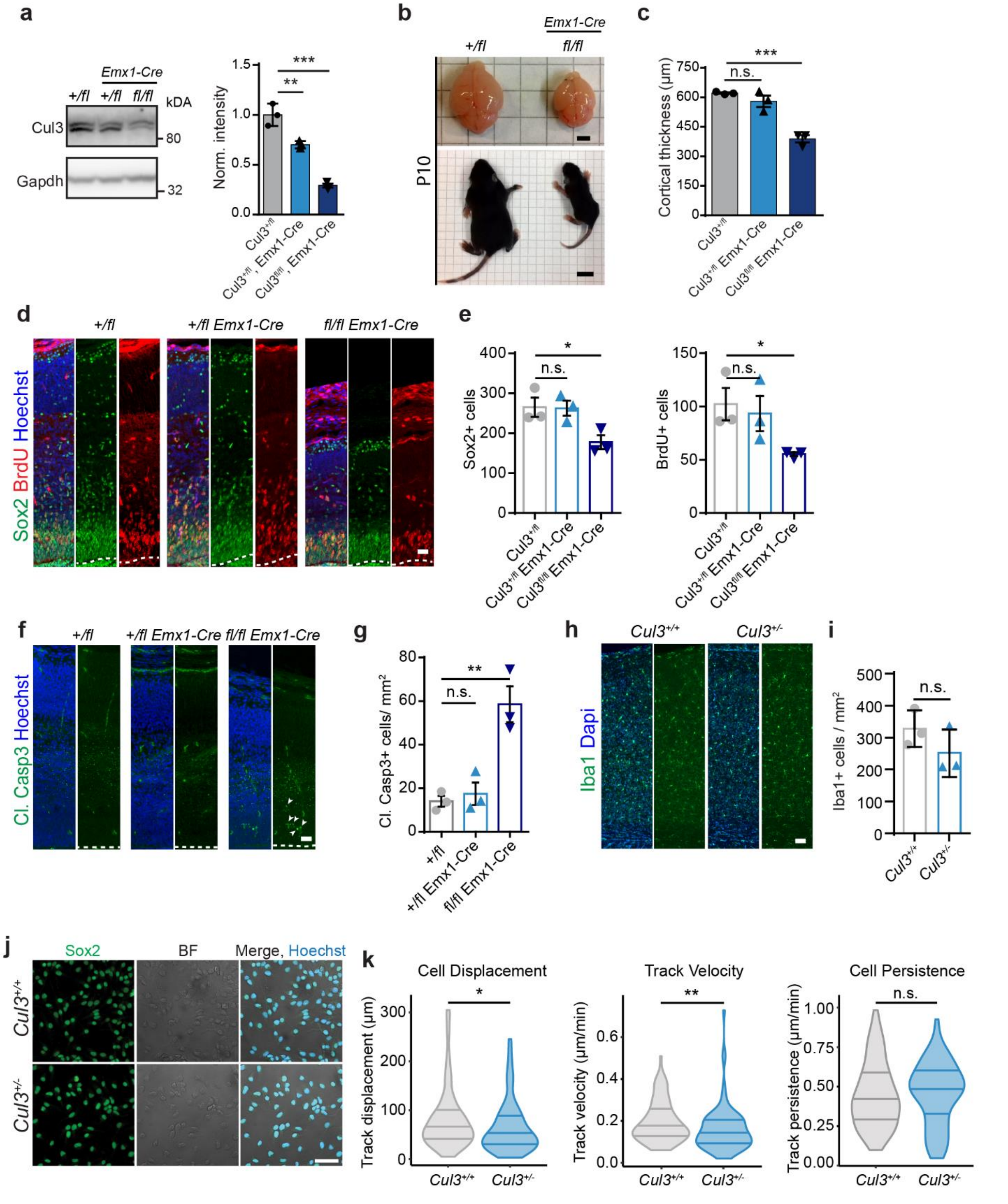

**Supplementary Figure 5. *Cul3* loss leads to reduced neuronal survival, neurogenesis and abnormal cell migration in mice. a,** Western blot and quantification of *Cul3*$^{+/fl}$, *Cul3*$^{+/fl}$ *Emx1-Cre* and *Cul3*$^{fl/fl}$ *Emx1-Cre* E16.5 brain lysates shows a strong reduction of Cul3 levels in conditional homozygous embryos (*n*= 3 littermate embryos per genotype; $^{**}P<0.01$, $^{***}P<0.001$; 1-way

ANOVA and Sidak's multiple comparisons test). **b**, Representative images of ten day-old *Cul3*$^{fl/fl}$ *Emx1-Cre* pups showing that mutant animals are much smaller than their *Cul3*$^{+/fl}$ control littermates. While hindbrain regions are comparable, forebrain structures are severely reduced in size in the conditional homozygous animals. **c**, Cortical thickness measured in Nissl stainings of coronal brain sections from *Cul3*$^{+/fl}$, *Cul3*$^{+/fl}$ *Emx1-Cre* and *Cul3*$^{fl/fl}$ *Emx1-Cre* newborn pups (P0), show severe cortical thinning in the latter (*n*= 3 pups per genotype; ***P<0.001, n.s. not significant; 1-way ANOVA and Sidak's multiple comparisons test). **d**, Representative images of E16.5 coronal brain sections, stained for the radial glia marker Sox2 and for BrdU incorporation (2 hour pulse) in *Cul3*$^{+/fl}$, *Cul3*$^{+/fl}$ *Emx1-Cre* and *Cul3*$^{fl/fl}$ *Emx1-Cre* embryos. **e**, Quantification of Sox2+ cells and BrdU+ cells reveals decreased numbers of cycling radial glia cells in the *Cul3*$^{fl/fl}$ *Emx1-Cre* developing forebrain (*n*= 3 littermates per genotype; *P<0.05; 1-way ANOVA and Sidak's multiple comparisons tests). **f**-**g**, Immunofluorescent staining for the apoptotic marker cleaved Caspase-3 shows increased cell death in the *Cul3*$^{fl/fl}$ *Emx1-Cre* E16.5 cortex (arrowheads: cl. Casp3+ cells) (*n*= 3 littermates per genotype; **P<0.01, n.s. not significant; 1-way ANOVA and Sidak's multiple comparisons tests). **h**-**i**, No difference in the number of Iba1+ microglia in the adult *Cul3*$^{+/-}$ cortex (*n*= 3 per genotype; n.s. not significant; unpaired two-tailed t-test). **j**, Representative pictures of *Cul3*$^{+/+}$ and *Cul3*$^{+/-}$ NPC preparations stained for Sox2. **k**, Cell tracks of NPCs detaching from neurosphere into embedding bovine collagen matrix imaged in a single plane. Cell displacement, cell velocity and migratory persistence were quantified in the imaging plane and compared between *Cul3*$^{+/+}$ and *Cul3*$^{+/-}$ cells (*n*(spheres)= 3 per genotype, *n*(cells)= 30 per replicate; *P<0.05; **P<0.01; Wilcoxon rank sum test). Data presented as mean ± SEM or violin plots with median and first and third quartiles. Scale bars: 2.5 mm in (b, top), 1 cm in (b, bottom), 25 µm in (d,f), 50 µm in (h) and 50 µm in (j). Detailed statistics can be found in Supplementary Table 1.

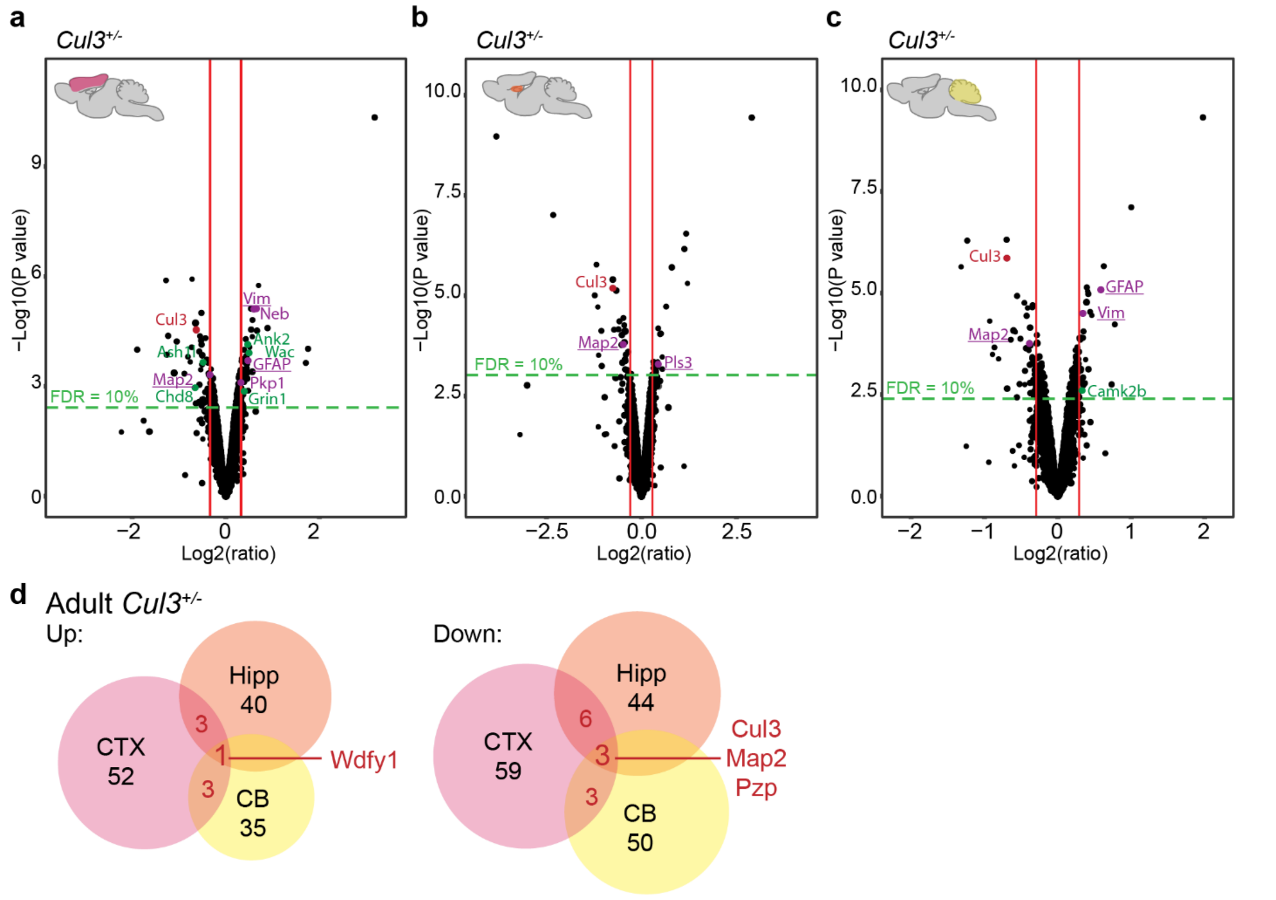


**Supplementary Figure 6. Minor protein composition alterations in the adult *Cul3*$^{+/-}$ brain. a**, Volcano plot of deregulated proteins at 10% FDR cut-off in the adult *Cul3*$^{+/-}$ cortex with 46 up- and 49 down-regulated protein groups (details in Supplementary Table 8). **b**, Volcano plot of deregulated proteins at 10% FDR cut-off in the adult *Cul3*$^{+/-}$ hippocampus with 19 up- and 28 down-regulated protein groups (details in Supplementary Table 9). **c**, Volcano plot of deregulated proteins at 10% FDR cut-off in the adult *Cul3*$^{+/-}$ cerebellum with 29 up- and 42 down-regulated protein groups (details in Supplementary Table 10). Purple: Cytoskeletal proteins, green: ASD-risk genes, red: Cul3 (a-c). **d**, Overlaps between up- and down-regulated protein groups in adult *Cul3*$^{+/-}$ brain tissues at 20% FDR (details in Supplementary Table 7).

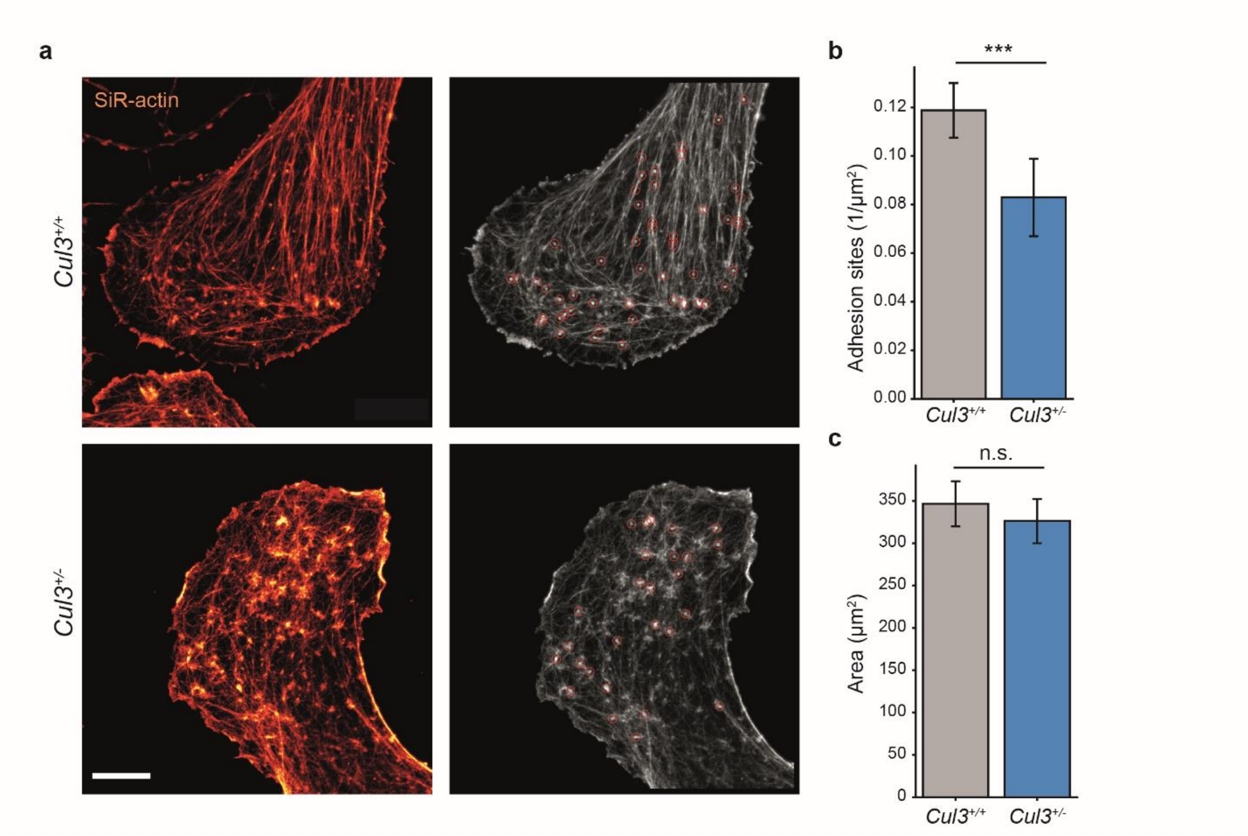


**Supplementary Figure 7. Decreased number of focal adhesions in *Cul3*$^{+/-}$ NPCs. a**, NPCs cultured on Poly-L-ornithine/Laminin were stained using SiR-actin (a) and leading edges of cell protrusions were imaged employing STED-microscopy (same images as in Fig. 6a). **b**, The number of bright SiR-actin puncta (red circles in a, right), putative focal adhesion sites, are reduced in *Cul3*$^{+/-}$ NPCs growth cones. Puncta were counted and normalized to the leading edge area (*n*(cells)= 43 per genotype from three independent NPCs preparations; ***P<0.001; two-tailed Welch's t-test). **c**, The growth cone areas were comparable between wild-type and mutant cells (*n*(cells)= 43 per genotype from three independent NPCs preparations, n.s. not significant, two-tailed Welch's t-test). Scale bar: 5 µm in (a). Detailed statistics can be found in Supplementary Table 1.

### Materials and Methods

#### Mice

We thank Dr. Jeffrey Singer (Portland State University, US) for providing us with the *Cul3*$^{flox}$ (*Cul3*$^{fl}$) conditional mouse line in which the exons 4 to 7 are flanked by loxP sites [46]. The *Cul3* constitutive knockout mouse line (*Cul3*$^{+/-}$) was obtained by mating *Cul3*$^{flox}$ mice with a CMV-Cre line (B6.C-Tg(CMV-cre)1Cgn/J) and back-crossings to C57BL/6J wild-type animals (Supplementary Fig. 1a). Genotyping for the *Cul3* knockout allele was performed using the following primers: *forward* GGAAACCTAAAGTTTTTATGCARG and *reverse* TTTGTCTGGACCAAATATGGCAGCCCAA ACC. The *Cul3 Emx1-Cre* conditional line was generated by crossing male *Cul3*$^{+/fl}$ mice with a *Emx1-Cre* expressing line (B6.129S2 (Emx1tm1cre)Krj/J). *Cul3*$^{fl/fl}$ *Emx1-Cre* mutant pups were sacrificed as soon as the phenotype was clearly detectable (strongly reduced size and very weak animals) within the first week of life, to comply with ethical requirements (3R principle). Embryonic time points were determined by plug checks, defining embryonic day (E) 0.5 as the morning after copulation. Animals were housed in groups of 3-4 animals per cage and kept on a 12 hour light/dark cycle (lights on at 7:00 am), with food and water available *ad libitum*. All animal protocols were approved by the Institutional Animal Care and Use Committee at IST Austria and the Bundesministerium für Bildung, Wissenschaft und Forschung, Austria, approval number: BMWFW-66.018/0009-WF/V/3b/.

#### Behavior

All behavioral tests were performed on adult, two- to five-month-old, sex-matched littermate animals during the light period. Animal cohort sizes ranged between 8 and 20 animals per genotype and sex. Male and female cohorts were initially analyzed separately, and data was only pooled in case wild-type male and wild-type female data was statistically comparable, i.e. not significantly different. Before testing, animals were habituated at least for one hour to the testing room, equipment was cleaned with 70% EtOH after each animal. Different behavioral tests in the same mouse cohort were separated by at least one day of break. Tests were performed starting with the least aversive task and ending with the most aversive, and either scored automatically or by an experimenter blind to the genotype.

***Hind limb clasping:*** During a 10 sec tail suspension period hind limb clasping severity (scores 0-1- no hindlimb clasping to 3- most severe phenotype) was assessed by the experimenter. The test was repeated 3 times per animal and the average score was calculated, scores between 0 and 1 are not considered as hindlimb clasping.

***Gait analysis:*** Gait properties were assessed by footprint analysis as described previously [47]. Briefly, fore- and hindpaws were colored with non-toxic dye and the animals allowed crossing a white sheet of paper, 70 cm in length, in a straight line. Stride, sway and stance length, as well as paw distance were measured as indicated in Fig. 1c by the experimenter.

***RotaRod:*** Performance on the accelerating mouse RotaRod (Ugo Basile S.R.L, Cat. No. 47650) was analyzed on 3 trials per day over three consecutive days, with an acceleration from 5 to 40 rpm over 5 min (300 sec). For each animal the latency to fall (sec) and maximum speed at the end of each trial (rpm) were automatically determined, sex-matched littermates were tested simultaneously to avoid confounding factors. Initial coordination was assessed by analyzing the first trial on the first day of training.

***Open field:*** As previously described [48], animals were allowed to freely explore a brightly lit arena (45 x 45 x 30 $cm^3$), made out of grey Plexiglas, over a 30 min time period. Locomotor activity (distance travelled and velocity) and center crossings were recorded by a video camera and analyzed using the EthoVision XT software (Noldus).

***Elevated plus maze (EPM):*** Mice were placed in the center of the EPM apparatus facing the open arm and left to explore the maze for 6 min. The time spent in the open and closed arms along with number of closed and open arm entries and distance travelled in each arm were determined using Ethovision XT video tracking system and software. The ratio of the time (in sec) spent on the open arm vs. the total time spent on the maze were calculated.

***Three chamber sociability test:*** Mice were tested for sociability and social novelty preference as described previously [47]. The testing apparatus was a rectangular clear Plexiglas three chambers box (60cm (L) x 40cm (W) x 20 cm (H)). The dividing walls had doorways allowing access to each chamber. Age and sex matched C57BL/6J mice were used as stranger mice and were habituated to placement inside the wire cage. Each test animal was first placed into the center chamber with open access to both left and right chamber, each chamber containing an empty round wire cage. The wire cage (12 cm height, 11 cm diameter) allows nose contact between mice but prevents fighting. After 10 min of habituation, during the social phase, an age-matched stranger was placed in the left chamber while a novel object was placed into the right chamber. The test animal was allowed to freely explore the social apparatus for 10 min. Subsequently, each mouse was tested in a second 10 min session to evaluate the preference for a novel stranger, which was placed inside the right wire cage. Number of nose contacts (< 5cm proximity) with the caged mouse, as well as the time spent in each chamber, was calculated. Analysis was done using the Noldus EthoVison XT.

***Olfaction habituation and dishabituation test (OHDH):*** The test was performed as previously described [49]. In brief, mice were presented with a sequence of non-social, i.e. water, almond (McCormick) and banana (McCormick) (diluted 1:100 in water), and social cotton swabs (Social A and B). Social odors were obtained by wiping the cotton swabs in a zick-zack manner through the bedding of two distinct cages, housing each three, to the test mice, sex- and age-matched wild-type mice. Each cotton odor swab was presented 3 x 2 min with a 1 min break. The time the tested mouse spent sniffing each cotton swab was determined by the experimenter using a stop swatch.

***Contextual fear conditioning (CFC) task***: As described previously [48], mice were subjected to the CFC task in three sessions, each distanced by 24 h: a training session (day 1) and two re-exposure sessions (day 2 and day 3). On day one, mice were subjected to a single fear conditioning training session of 5 min, in which they learned to associate the conditioned stimulus (CS: context) to the unconditioned stimulus (US: a foot-shock). To this end, each mouse was placed in a fear-conditioning chamber (18 x 18 $cm^2$, Noldus) with an electrified grid-floor. After 120 sec of free exploration, the mouse was subjected three times to a foot-shock (0.5 mA; 2 sec) delivered through the grid-floor, every 1 min. Mice remained in the conditioning chamber for 1 min after the last shock delivery. On days two and three, each mouse was placed back in the conditioned chamber for 10 and 5 min, respectively, without delivering the electrical foot-shock to test their memory retention and memory extinction. Behavior during all experimental sessions was recorded by a video camera mounted above the ceiling of the cage and connected to a computer equipped with the Ethovision XT software (Noldus). Percentage of time spent freezing (absence of all but respiratory movements for at least 3 sec) was scored to assess emotional reactivity during training (day 1) and fear memory during retention test (the first 3 min of day 2 session) and extinction test (first 3 min of day 3 session). All behavioral parameters were scored by an experimenter blind to the animal experimental condition. Pairs of control-mutant littermates were randomly tested in the morning and in the afternoon to control for circadian rhythm.

**Immunofluorescence; BrdU labeling and imaging**

***Immunofluorescent staining in adults:***

Adult male littermate mice were deeply anesthetized and transcardially perfused with 4% paraformaldehyde (PFA). Brains were dissected, postfixed in 4% PFA, dehydrated in 30% sucrose and sliced at 40 µm on a sliding VT1200S vibratome (Leica Microsystems). Stainings on adult brains were performed on floating sections without antigen retrieval. In brief, sections were washed in 1x PBS and incubated overnight on a horizontal shaker at 4°C in primary antibody solution (14-16 hours). Primary antibodies were diluted in 0.3% Triton X-100 and 2-5% donkey serum. On the next day, the sections were washed and incubated with a species-specific secondary antibody for 2 hours at 4°C. Nuclear counterstain was performed for 10 min with 300 nM DAPI (Life Technologies) in 1x PBS before mounting in DAKO fluorescent mounting medium. To examine cortical layering, the thickness of Cux1-positive cell layer and the Ctip2-positive cell layer was measured at three defined points of each cortical hemisphere (*n*= 3 littermate animals per genotype, at least 4 images/animal). For interneuron, microglia and oligodendrocyte counting, positive-stained cells were counted within the somatosensory cortex and normalized to the area used (*n*= 3 littermate animals per genotype, at least 5 images/animal). Adult cortical sections were stained with the following primary antibodies: anti-Cux1 (Santa Cruz, sc-13024, 1:200), anti-Ctip2 (Abcam, ab18465, 1:500), anti-Parvalbumin (1:500, Chemicon MAB1572), anti-Iba1 (1:500, Wako 019 19741), anti-GFAP (1:200, Cell Signaling 12389P).

***Immunofluorescent staining in embryos and newborn pups (P0/P1):***

Mice were killed for analysis at E14.5 and E16.5 and P0 by decapitation. Heads were dropfixed in 4% PFA overnight, dehydrated in 30% sucrose, embedded in O.C.T. (Tissue Tek) and 18 µm sections were prepared on a Microm HM560 cryostat (Thermo Scientific). For assessment of the cellular composition of the embryonic cortex by immunofluorescent stainings, antigen retrieval was performed using 1x DAKO antigen retrieval solution (s1699) and immunofluorescent staining was performed as outlined above. The following primary antibodies were used: anti-Cux1 (Santa Cruz, sc-13024, 1:500), anti-Ctip2 (Abcam, ab18465, 1:500), anti-cleaved Caspase3 (Cell Signaling, 9661, 1:300), anti-Sox2 (Millipore, AB5603, 1:200). Analysis of Cux1/Ctip2 cell distributions at P0 was performed by counting the relative number of Cux+ and Ctip2+ cells in each of 10 bins (bin height adjusted to cortical thickness, bin width 200 µm) in the cortex, normalized to the total number of positive cells in the respective region (*n*= at least 3 littermate mice per genotype, at least 4 images/animal).

***BrdU based birthdate labeling for migration and proliferation analysis:*** For **cell cycle analysis,** pregnant mice were injected with 0.1 mg/g bromodeoxyuridine (BrdU) at E14.5 and E16.5 and sacrificed 2 hours later; embryos were decapitated and processed for immunostaining as described above. Cells in S-phase that incorporated BrdU were detected using anti-BrdU (BioRad, MCA2060T, 1:500). The number of BrdU+ cells was manually counted in cortical regions of 200 µm width in blinded images (*n*= 3 littermate animals per genotype, at least 5 images/animal). For **migration analysis,** pregnant females were injected with BrdU at E16.5, as described elsewhere [50]. Upon delivery (P0), pups were decapitated and tissue was processed for immunostaining as outlined above. For migration analysis, sections were stained with anti-BrdU (BioRad, MCA2060T, 1:500), and anti-Ctip2 (Abcam, ab18465, 1:500) antibodies. The P0 cortex was divided into ten bins of the same size and the relative number of BrdU+ nuclei per bin counted manually in blinded images (*n*= at least 3 littermate mice per genotype, at least 5 images/animal).

***Imaging:*** Images from immunofluorescent stainings were acquired on a Zeiss LSM800 inverted confocal microscope, background corrected and adjusted for contrast and brightness, as well as analyzed in Fiji [51] using the cell counter plugin.

***Nissl staining and Golgi staining:***

***Nissl staining:*** For Nissl staining, brains from perfused adult animals were post-fixed in 4% PFA overnight, dehydrated and paraffin embedded. Sagittal and coronal sections were cut on a Microtome HM 355 at 10 µm thickness. For Nissl stainings in newborn mice, pups were decapitated at P0, brains dissected, dropfixed in 4% PFA, dehydrated in sucrose, embedded in O.C.T and cut at 18 µm on a cryostat. Nissl staining with 1% Cresyl Violet solution (Cresyl Violet Acetate, Sigma, Cat.No C 5042) was performed upon clearance of paraffin slices with RotiHistol

(Carl Roth) for 10 min and rehydration of sections (absolute EtOH to water: 96%, 90%, 70%, 50%, 30%, water, 3-5 min each), or 3x 5 min washes in 1x PBS to remove the OCT. Nissl stainings of adult and P0/P1 brains were captured using a Olympus Slide scanner VS120 and analyzed using Fiji.

***Golgi staining and analysis:*** Golgi-Cox staining was performed according to protocol using the FD Rapid GolgiStain Kit$^{TM}$ (FD Neurotechnologies). After three weeks of Golgi impregnation, brains were cut coronally (120 µm) using a Leica Vibratome (Leica VT 1200S) and mounted onto 1% gelatin-coated slides. Slides were then dehydrated through graded ethanol steps, cleared with RotiHistol (Carl Roth) and mounted with DPX mounting medium on coverslips (#1.5). To quantitatively analyze pyramidal neurons in Golgi-stained slides, impregnated pyramidal cells (8-10 neurons per brains, *n*= 3 littermate brains per genotype) of layer 2/3 in the somatosensory cortex were selected and imaged with a Nikon Eclipse Ti2 using a 40x magnification. For analysis, single pyramidal neurons were manually reconstructed using Imaris analysis software (version 9.3.1). The average filament area, filament length and Sholl intersections were analyzed using the same software. Spine counting was performed using Fiji, spines that started from 100 µm distance of the apical dendrite were counted within a 100 µm segment.

**LC-MS/MS/MS whole proteome analysis**

***Samples:*** Adult male *Cul3*$^{+/-}$ and *Cul3*$^{+/+}$ wild-type littermate animals (*n*= 5 littermate mice/genotype) were deeply anesthetized and transcardially perfused with 15 ml of ice-cold 0.9% NaCl to clear the brain from blood. The cortex, hippocampus and cerebellum were rapidly dissected on ice, snap-frozen in liquid nitrogen and stored at -80° until protein extraction. Embryonic E16.5 forebrain tissue (developing cortex and hippocampus; male embryos *n*(constitutive)= 5 mice/genotype, *n*(Emx1Cre conditional)= 3 mice/genotype) was dissected on ice, meninges were removed and snap-frozen in liquid nitrogen and stored at -80° until protein extraction. Tissues were homogenized 1:5 (w:v) in modified RIPA buffer (50 mM Tris-HCl pH 7.5, 150 mM NaCl, 1% NP40, 0.5% Sodium deoxycholate, 0.1% SDS, 1mM EDTA, 10mM NaF) and freshly added protease and phosphatase inhibitors (Roche 04 693 159 001 and 04 906 837 001) and lysed for 30-45 min on ice, while occasionally being vortexed gently. Each sample was sonicated twice at 180 W in an ice-cold water bath and centrifuged at 10.000 rpm for 20 min at 4°C. Lysates were quantified using the Pierce™ BCA Protein Assay Kit (Thermo Fisher, Cat. no. 23225).

***TMT Labeling and High pH reversed-phase chromatography:*** Aliquots of 100 µg of each sample were digested with trypsin (2.5 µg trypsin per 100 µg protein; 37°C, overnight), labeled with Tandem Mass Tag (TMT) 11plex reagents according to the manufacturer's protocol (Thermo Fisher Scientific, Loughborough, LE11 5RG, UK) and pooled. For the adult dataset, where the number of samples exceeded the number of available TMT channel, one combined TMT sample was generated for each tissue. Pooled samples were evaporated to dryness, re-dissolved in 5% formic acid and then desalted using a SepPak cartridge according to the manufacturer's instructions (Waters, Milford, Massachusetts, USA). Eluate from the SepPak cartridge was again evaporated to dryness and re-dissolved in buffer A (20 mM ammonium hydroxide, pH 10) prior to fractionation by high pH reversed-phase chromatography using an Ultimate 3000 liquid chromatography system (Thermo Scientific). In brief, the sample was loaded onto an XBridge BEH C18 Column (130Å, 3.5 µm, 2.1 mm X 150 mm, Waters, UK) in buffer A and peptides eluted with an increasing gradient of buffer B (20 mM Ammonium Hydroxide in 90% acetonitrile, pH 10) from 0-95% over 60 min. The resulting fractions were evaporated to dryness and redissolved in 1% formic acid prior to analysis by nano-LC MSMS using an Orbitrap Fusion Lumos mass spectrometer (Thermo Scientific).

***Nano-LC Mass Spectrometry:*** High pH RP fractions were further fractionated using an Ultimate 3000 nano-LC system in line with an Orbitrap Fusion Lumos mass spectrometer

(Thermo Scientific). In brief, peptides in 1% (V/V) formic acid were injected onto an Acclaim PepMap C18 nano-trap column (Thermo Scientific). After washing with 0.5% (V/V) acetonitrile 0.1% (V/V) formic acid, peptides were resolved on a 250 mm × 75 μm Acclaim PepMap C18 reverse phase analytical column (Thermo Scientific) over a 150 min organic gradient, using 7 gradient segments (1-6% solvent B over 1min., 6-15% B over 58min., 15-32%B over 58min., 32-40%B over 5min., 40-90%B over 1min., held at 90%B for 6min and then reduced to 1%B over 1min.) with a flow rate of 300 nl $min^{-1}$. Solvent A was 0.1% formic acid and Solvent B was aqueous 80% acetonitrile in 0.1% formic acid. Peptides were ionized by nano-electrospray ionization at 2.0 kV using a stainless steel emitter with an internal diameter of 30 μm (Thermo Scientific) and a capillary temperature of 275°C.

All spectra were acquired using an Orbitrap Fusion Lumos mass spectrometer controlled by Xcalibur 4.1 software (Thermo Scientific) and operated in data-dependent acquisition mode using an SPS-MS3 workflow. FTMS1 spectra were collected at a resolution of 120,000, with an automatic gain control (AGC) target of 200,000 and a max injection time of 50 ms. Precursors were filtered with an intensity threshold of 5,000, according to charge state (to include charge states 2-7) and with monoisotopic peak determination set to Peptide. Previously interrogated precursors were excluded using a dynamic window (60s +/-10ppm). The MS2 precursors were isolated with a quadrupole isolation window of 0.7 m/z. ITMS2 spectra were collected with an AGC target of 10,000, max injection time of 70 ms and CID collision energy of 35%. For FTMS3 analysis, the Orbitrap was operated at 50,000 resolution with an AGC target of 50,000 and a max injection time of 105 ms. Precursors were fragmented by high energy collision dissociation (HCD) at a normalized collision energy of 60% to ensure maximal TMT reporter ion yield. Synchronous Precursor Selection (SPS) was enabled to include up to 5 MS2 fragment ions in the FTMS3 scan.

***Peptides Identification and TMT Reporter Quantitation:*** Acquired raw data files were processed and quantified using Proteome Discoverer software v2.1 (Thermo Scientific) and searched against the UniProt *Mus musculus* database (downloaded November 2018: 81925 sequences) using the SEQUEST algorithm. The raw files from the embryonic and adult samples were processed in two separate batches. For each, peptide precursor mass tolerance was set at 10 ppm, and MS/MS tolerance was set at 0.6 Da. Search criteria included oxidation of methionine (+15.995) and phosphorylation of serine, threonine or tyrosine (+79.966) as variable peptide modifications and carbamidomethylation of cysteine (+57.021) and the addition of the TMT mass tag (+229.163) to peptide N-termini and lysine as fixed modifications. Acetylation (+42.011) and Met-loss + Acetylation (-89.030) were included as possible modifications to the protein N-terminus. Searches were performed with full tryptic digestion and a maximum of 2 missed cleavages were allowed. The reverse database search option was enabled and all data was filtered to satisfy false discovery rate (FDR) of 5%.

***Statistical Data Analysis:*** PSMs tables exported from Proteome discoverer were reprocessed in R using in house scripts. PSMs reporter intensities were scaled by precursor integrated peak intensities and normalized across TMT channels. The resulting peptidoform expression matrix was then renormalized across samples using, successively, SVN normalization followed by the Levenberg-Marquardt procedure. In addition, the combined embryonic dataset was batch-corrected against the litter effect using the ComBat function from the SVA package (batch correction was skipped for the adult dataset as PCA analysis revealed only minor litter effect within tissues, and correcting for TMT batch would have removed tissue-specific variation, since each TMT sample was tissue specific). Peptidoforms were assembled into protein groups, and protein groups-level expression values calculated from those of individual peptidoforms, excluding phosphorylated peptides and their unmodified counterpart form. Average expression values and ratios were calculated for each condition, and moderated t-test performed using the limma package. P-value significance thresholds were calculated for pre-agreed 10, 20 and 30 % false discovery rate levels (Benjamini Hochberg procedure). In addition, regardless of P value, ratios were also not considered if the absolute value of their base-2 logarithm did not exceed a threshold calculated as excluding all but the 5% most extreme ratios between individual controls. For functional annotation clustering, proteins found significant for a particular condition were mapped to GO terms using the DAVID Bioinformatics Resources 6.8 online tool [52,53](done for proteins at FDR 10 and 20% FDR, GO- terms filtered for p< 0.01, Benjamini adjusted p-values are reported for significantly enriched terms). Predicted ASD-associated proteins were obtained from the SFARI gene database (https://gene.sfari.org/database/gene-scoring/) and a hyper-geometric test performed to establish significance.

**Western blots**

Littermate embryonic and adult animals (at least *n*= 3 per genotype) were decapitated, the brain was dissected on ice, snap frozen in liquid nitrogen and stored at -80°C until protein extraction. Tissues were homogenized in ice-cold RIPA buffer (50 mM Tris-HCl pH 7.5, 150 mM NaCl, 1% NP40, 0.5% Sodium deoxycholate, 0.1% SDS, 1mM EDTA, 10 mM NaF) and freshly added protease inhibitors (Roche), lysed for 45 min on ice and centrifuged at 10.000 rpm for 20 min at 4°C. Lysates were quantified using the Pierce™ BCA Protein Assay Kit (Thermo Fisher, Cat. no. 23225).

For Western Blots, 25-50 µg of proteins were mixed with 6X Laemmli buffer (375 mM Tris pH=6.8, 12% SDS, 60% glycerol, 600 mM DTT, 0.06% bromophenol blue), heat-denatured at 95°C and separated on 8-10% SDS-PAGE gels in running buffer. Proteins were transferred to a PVDF membrane (Merck) using transfer buffer in a Western blotting apparatus (Bio-Rad) for 2 h at 4°C with 300 mA constant current. The membranes were then blocked with 5% milk in 1x TBST for 1h at room temperature and incubated with primary antibodies overnight at 4°C. Secondary anti-IgG antibody coupled to horseradish peroxidase (HRP) was detected using a Pierce™ enhanced chemiluminescent substrate (ThermoFisher) on a GE Healthcare Amersham machine. The following primary antibodies were used: anti-Cul3 (1:800, Cell Signaling #2759), anti-Gapdh (1:1000, Merck ABS16), anti-Pls3 (T-Plastin 1:500, Thermo Fisher PA5-27883), anti-Pls1 (1:500, Novus Biologicals H00005357-M04), anti-alpha internexin (1:5000, Abcam ab40758), anti-Nischarin B3 (1:200, Santa Cruz sc-365364). Secondary antibodies used: donkey anti-rabbit IgG (1:5000, Amersham NA934) and goat anti-mouse IgG (1:10000, Pierce 31432).

**RNA isolation and quantitative real time PCR (qRT-PCR) analysis**

Tissue from E12.5, E14.5, E16.5, P1, P21 and adult C57BL6/J brains was used for wild-type expression analysis. Tissue from *Cul3*$^{+/-}$ and wild-type E16.5 embryos was used to validate up-regulated proteins identified through proteomic analyses. For RNA extraction 700 µL Trizol (Invitrogen) and 140 µL chloroform (Sigma) was used for homogenization, followed by centrifugation at 12,000 x *g* for 15 min at 4°C. The upper aqueous phase was transferred to a new tube and 1.5 volumes of 100% ethanol (EtOH) were added. RNA was extracted using Zymo-Spin™ IC columns (Zymo Research). In short, the aqueous phase/ethanol mixture was loaded onto the column, washed with 400 µl 70% EtOH before being treated with RQ1 DNAseI (Promega, 5 µl + 5 µl reaction buffer + 40 µl 70% EtOH) for 15 min at RT. After two washes with 70% EtOH the sample was eluted from the column with DEPC-treated $H_2O$. RNA concentration was measured by using the NanoDrop spectrophotometer (Thermo Scientific). 500 ng of RNA were used for cDNA preparation with the RevertAid First Strand cDNA Synthesis Kit (Thermo Fisher). cDNA was diluted 1:3 and used for qPCR analysis with Lightcycler 480 Sybr green master mix (Roche) on a Real-Time PCR Roche Lightcycler 480 machine. Samples were run in duplicates or triplicates with the following intron-spanning qPCR primers for target genes: *Cul3 forward:* AAGGTGGTGGAGAGGGAACT and *reverse* TCAAACCATTTGGCACACGAC; *Pls3 forward* TCTAGAAGGGGAAACTCGGG and *reverse* GGATCACCAGAGCATCCTGC; *Pls1* forward CCATGCCTACACAAGCCTGA and *reverse* GCGTCTGCAAGGTCACTGTA; *INA* *forward* CCAGGCACGTACCATTGAGAT and *reverse* CAATGCTGTCCTGGTAGCCG. As housekeeping genes *Pgk1* (*forward* AAAGTCAGCCATGTGAGCACT and *reverse* ACTTAGGAGCACAGGAACCAAA) and *Gapdh* (*forward* AACGGGAAGCTCACTGGCAT and *reverse* GCTTCACCACCTTCTTGATG) were used. ΔCq expression levels (relative mRNA) were calculated upon normalization to housekeeper genes and plotted.

**Region/Cell-type specific expression of *Cul3***

Brain-region specific RNA-seq data (as RPKM) were downloaded from the BrainSpan Atlas of the Developing Human Brain (http://www.brainspan.org/static/download.html). For the developmental trajectory, prefrontal cortex samples (DFC, MFC and VFC) were selected and the expression of *CUL3* was plotted for time points up to 1 year of age. For the brain region specific expression adult samples (starting at 18 years of age) were grouped according to region. After selection, the data was plotted as mean across samples with individual dots representing individual samples.

Cell-type specific data from scRNA-seq experiments and aggregated by cluster (as log2(CPM+1)) was downloaded from the Allen Cell Types Database (https://portal.brain-map.org/atlases-and-data/rnaseq, mouse: “Cell Diversity in the Mouse Cortex and Hippocampus”, human: “Cell Diversity in the Human Cortex”). Clusters were grouped according to broad cell type based on description and hierarchical relationship of the clusters and data was plotted as mean across clusters with individual dots representing individual clusters of the same broad cell type. Some clusters of rare cell types were omitted from the mouse data to simplify the plot.

**Electrophysiology**

Brain slices were obtained from *Cul3*$^{+/-}$ and wild-type male littermates. Acute coronal slices (300 µm) were prepared from primary somatosensory cortex. Animals were decapitated under isoflurane anesthesia and whole brains were rapidly removed from the skull and sectioned using a VT 1200S vibratome (Leica Microsystems) in ice-cold cutting solution, containing (mM): 93 NMDG, 2.5 KCl, 1.2 NaH2PO4, 30 NaHCO3, 20 HEPES, 25 glucose, 5 sodium ascorbate, 2 thiourea, 3 sodium pyruvate, 10 MgCl2, 0.5 CaCl2 (320 mOsm, 7.2-7.4 pH). Slices were recovered at 32˚C for 12 min in the same solution and then allowed to recover at room temperature for at least 1 hour in regular artificial cerebrospinal fluid (ACSF), containing (mM): 125 NaCl, 2.5 KCl, 1.25 NaH2PO4, 25 NaHCO3, 25 glucose, 1 MgCl2 and 2 CaCl2 (320 mOsm, 7.2–7.4 pH). The ACSF was continuously oxygenated with 95% $O_2$ and 5% $CO_2$ to maintain the physiological pH. Slices were visualized under infrared-differential interference contrast (IR-DIC) using a BX-51WI microscope (Olympus) with a QIClickTM charge-coupled device camera (Q Imaging Inc, Surrey, BC, Canada).

Patch pipettes (3–5 MΩ; World Precision Instruments) were pulled on a P-1000 puller (Sutter Instruments) and filled with the intracellular recording solution, containing (mM): 115 cesium methanesulphonate, 8 NaCl, 10 HEPES, 0.3 EGTA, 10 Cs4BAPTA, 4 MgATP, 0.3 NaGTP and 0.2% biocytin. Internal pH was adjusted to 7.3 with CsOH and osmolarity adjusted to 295 mOsm with sucrose. Spontaneous postsynaptic currents were recorded from pyramidal neurons in layer 2/3. Excitatory currents (sEPSC) were recorded at holding potential -70 mV, while inhibitory currents (sIPSC) at +10 mV. Signals were filtered at 2 kHz, digitized at 10 kHz and acquired using a MultiClamp 700B amplifier and a Digidata 1550A. Recorded signals were low-pass filtered at 1 kHz and analyzed using Clampfit 10 software (Molecular Devices). Both excitatory and inhibitory synaptic currents were identified by a template created for each neuron using 50-100 single events for each trace. All events recognized through the template were visualized, identified and accepted by manual analysis. Cumulative distributions for single neurons and recording conditions were obtained by pooling together 300–400 single synaptic currents.

**Generation and culture of neural progenitor cells**
Neural progenitor cells (NPCs) were generated from E13.5 mouse cortices. Briefly, cortical tissues were dissected in L15 medium (Sigma cat. nr. L5520). The tissue was dissociated using Accutase (Sigma cat. nr. A6964) for 5 min at 37°C, pelleted in basal media, re-suspended in complete media and cells plated in uncoated dishes. The next day, cells were dissociated using Accutase, pelleted and re-suspended in complete media and pooled according to their genotype (*Cul3*$^{+/+}$ and *Cul3*$^{+/-}$ NPC) for primary neurosphere formation. Two independent NPC batches were prepared and analyzed. For adherent NPC cultures, dishes were coated with 10 µg/ml Poly-L-ornithine (PLO) (Sigma cat. nr. P3655) and Laminin (3mg/ml Sigma cat. nr. L2020) and approximately 15 000 cells/cm$^2$ cells were plated. 1x10$^6$ cells were frozen per freezing vial using a freezing medium containing basal medium, DMSO and fetal bovine serum (FBS) (8:1:1). Composition of media: Basal medium: 1x DMEM/F12 (Gibco cat. nr. 32500-350), 2 mM Glutamax (Gibco cat. nr. 35050061), 15 mM HEPES (Sigma cat. nr. H0887), 2% D-Glucose (Sigma cat. nr. G879), Sodium bicarbonate (Sigma cat. no. S8761), 100X Penicillin/ Streptomycin (Sigma cat. nr. P4333) in sterile water. Complete medium: Basal media plus freshly added 20ng/ ml human recombinant EGF (R&D cat. nr. 236-EG) and 10ng/ mlhuman recombinant bFGF (R&D cat. nr. 233-FB-025), B27™ Supplement minus VitA (Gibco cat. nr. 12587010).

***In vitro* migration assay and live cell imaging**
For in *vitro* migration assays, NPCs were dissociated using Accutase for 5 min at 37°C, pelleted in basal media and re-suspended in complete media. Two thousand cells were seeded per well in U-bottom 96-well ultralow attachment plates (Corning), assembled into neurospheres overnight (14-16 hours). Healthy neurospheres (defined by smooth and bright surface) were embedded under glass cover slips in 30 µl of hanging drops of 3D collagen scaffold with a final concentration of 1.7 mg/ml (obtained by mixing bovine collagen (PureCol, Advanced BioMatrix, USA) in 1X minimum essential medium eagle and 0.4 % sodium bicarbonate (both Sigma-Aldrich, USA). Collagen was let to polymerize at 37°C with 5% $CO_2$ and humidity for 1 hour. Next, dishes were flipped around and a second layer of collagen was added on top and let polymerize for another 1 hour. Devices were covered with complete media and imaged using a 10x objective on a brightfield inverted microscope at 37°C with 5% $CO_2$ for 72 hours every 10 min. Calibrated time course TIFF stacks were created in Fiji. Individual cells were tracked manually by an experimenter blinded to the genotype employing the Fiji plugin TrackMate [54]. Cells were chosen randomly from all sites of the sphere at the moment of their detachment and tracked until either re-joining the growing spheres or until the end of the recording. TrackMate data was then imported into R (3.6.2) environment and analyzed. The track velocity (track displacement / track duration) of detaching cells, their average instantaneous speed (average of all displacements per frame duration), track path length (sum of all displacements between frames) and persistence (track displacement / track path length) were quantified by TrackMate and with R. Finally, results were plotted with the help of ggplot2 (3.2.1), ggpubr (0.2.4) packages.

Alternatively, an adapted neurosphere migration assay was used [55]. Spheres were embedded centrally in 40 µl of 5 mg/ml matrigel (Corning cat. nr. 356234) diluted in ice-cold basal media on PLO/Laminin coated clear bottom 96-well imaging plates (Corning cat. nr. 3603), one sphere per well. After 30 min polymerization of the matrigel at 37°C, complete media was added to the wells and images were captured using a 4x objective on a brightfield inverted microscope, to determine initial sphere sizes. Neurospheres too close to the well-border were excluded from analysis. 22 and 46 hours after embedding, radial outgrowth of NPCs from the sphere was imaged using a 4x objective on a brightfield inverted microscope and the distance (radius) from the center of the neurospheres was measured in Fiji and normalized to the initial sphere radius.

**STED microscopy**

***Dual color labeling of neural progenitor cells (NPCs) for actin and tubulin*:** NPCs were seeded at a confluency of 70% on #1.5H glass coverslips coated with PLO/Laminin (Marienfeld, Lauda-Königshofen, Germany). After ~12 – 14 hours they were fixed with 4% paraformaldehyde in PBS for 15 min at room temperature (RT) followed by 3 x 2 min in PBS. Permeabilization was performed with PBS + 0.25% Triton X-100 (Sigma-Aldrich) for 10 min at RT followed by 3 x 2 min in PBS. Cells were blocked with 2% BSA (AppliChem GmbH, Darmstadt, Germany) in PBS for 30 min at RT and then incubated with anti-α-tubulin antibody (T6074, Sigma-Aldrich; 1:1000 in blocking solution) for 1 hour at RT. Samples were washed 3 x 3 min in PBS at RT. Incubation with secondary antibody and simultaneously with SiR-actin was done in blocking solution at RT for 1 hour (SiR-actin, SC001, Spirochrome, Switzerland, 1:500; goat anti-mouse IgG, conjugated to Alexa Fluor 594, Invitrogen A11005, 1:500). Samples were washed 3 x 3 min in PBS at RT and then incubated with DAPI (D9542, Sigma-Aldrich; 1:5000) in PBS at RT for 5 min. Samples were washed 3 x 2 min in PBS, then mounted with Dako Fluorescence Mounting Medium (S3023, Agilent Technologies).

***STED microscopy:*** STED imaging was performed on an Abberior Instruments Expert Line STED microscope. STED wavelength was 775 nm and excitation was performed at ~560 nm and ~640 nm. A 100x/1.4 NA oil immersion objective (Olympus, UPLSAPO 100XO) was used. STED pulses had a duration of ~1 ns and time gating was applied throughout. Resolution was increased in the xy-direction with a “doughnut” shaped beam and in the z-direction with an additional z-STED pattern according to the power ratio given below. Power levels are given as power at the back aperture of the objective lens with an estimated uncertainty of ~15%.

***Imaging parameters:***

***Fig. 6a top Cul3$^{+/+}$:*** STED: pixel size 30 nm x 30 nm; pinhole size 1 Airy Unit; Excitation laser powers: 560 nm: 5.4 µW; 640 nm: 2.9 µW; STED: 90 mW with lateral/axial STED power ratio 75/25. Pixel dwell time: 15 µs with 2 scans per line for 560 nm excitation and 4 scans per line for 640 nm excitation.

***Fig. 6a bottom Cul3$^{+/-}$:*** STED: pixel size 30 nm x 30 nm; pinhole size 1 Airy Unit; Excitation laser powers: 560 nm: 2.3 µW; 640 nm: 6.2 µW; STED: 90 mW with lateral/axial STED power ratio 75/25. Pixel dwell time: 15 µs with 2 scans per line for 560 nm excitation and 4 scans per line for 640 nm excitation.

***Fig. 6d top Cul3$^{+/+}$:*** STED: pixel size 30 nm x 30 nm; pinhole size 1 Airy Unit; Excitation laser powers: 560nm: 5.4 µW; 640 nm: 2.9 µW; STED: 90 mW with lateral/axial STED power ratio 75/25. Pixel dwell time: 15 µs with 2 scans per line for 560 nm excitation and 4 scans per line for 640nm excitation.

***Fig. 6d bottom Cul3$^{+/-}$:*** STED: pixel size 30 nm x 30 nm; pinhole size 1 Airy Unit; Excitation laser powers: 560 nm: 2.3 µW; 640 nm: 6.2 µW; STED: 90 mW with lateral/axial STED power ratio 75/25. Pixel dwell time: 15 µs with 2 scans per line for 560 nm excitation and 4 scans per line for 640nm excitation.

**Orientation analysis**

The orientation of actin and tubulin fibers was analyzed in custom-written Python routines using the scikit-image library [56] (ver. 0.16.2). We extracted the orientation distributions based on the structure tensor. The structure tensor summarizes local orientations and their coherence (degree of anisotropy) at each image location [57]. For each image of size 30 µm x 30 µm with pixel size 30 nm x 30 nm (1000 pixel$^2$), we first normalized the raw photon counts from 16-bit integers to values in the range [0, 1] by subtracting the minimum gray value, followed by dividing with the maximum. From the structure tensor computed at Gaussian scale σ=60 nm (2 pixel), we extracted local orientations, coherency, and energy (i.e. image gradient magnitude). Coherence yields values of 1 when the local structure is totally aligned, and 0 when there is no preferred direction. The energy measures the magnitude of the local structure (gradient). The resulting orientations were visualized in the hue, saturation, and brightness (HSB) color-space, where we set hue as the orientation angle, saturation as coherency and brightness the normalized gray-value of the source image. The distribution of orientations was built as the histogram of orientations weighted by the corresponding coherency. Image locations with a coherence or energy less than 10% of their maximum value were excluded from the histogram. Each orientation histogram was aligned to the corresponding dominant orientation angle computed from the tubulin channel. The dominant orientation is computed as the average orientation angle inside the cell. Each cell was segmented manually prior to analysis.

After alignment, we averaged all orientation histograms per group (*Cul3*$^{+/+}$, *n*= 43; *Cul3*$^{+/-}$, *n*= 43, from three independent NPC preparations respectively) and computed 95% confidence intervals per orientation angle. To test statistical significance, we measured the spread of each orientation distribution as the standard deviation of angles weighted by their occurrences. Reported p-values were computed using a two-tailed Welch's t-test using the scipy library (ver.1.3.0).

**Adhesion site analysis**

Adhesion sites in the actin channel were analyzed in custom-written Python routines using the scikit-image library. Each cell was segmented manually prior to analysis. For each image of size 30 µm x 30 µm with pixel size 30 nm x 30 nm (1000 pixel²), we first normalized the raw photon counts from 16-bit integers to values in the range [0, 1] by subtracting the minimum gray value, followed by dividing with the 99.9$^{th}$ percentile. We counted the number of adhesion sites per cell as the number of local maxima in the scale-space of a series of Laplacian of Gaussian (LoG) filters (https://doi.org/10.1007/BF01469346). A LoG filter with scale σ is a high-pass filter with a strong response at bright, blob-like structures of radius $\sqrt{2\sigma}$. We computed a series of 5 LoG filters for σ = 3, 4, 5, 6, and 7 pixel (90, 120, 150, 180, and 210 nm). A local maximum was considered an adhesion site if it exceeded a threshold of 0.25 in the LoG filter response. To exclude detection at the cell boundary we removed adhesion sites closer than 1 µm from the manually segmented cell boundary. To compute the density of adhesion sites, we divided the number of detected adhesion sites by the area of the cell. To test statistical significance we used a two-tailed Welch's t-test using the scipy library (ver 1.3.0).

**Tamoxifen induced *Cul3* deletion**

To induce *Cul3* deletion in adult animals double transgenic mice, *Cul3*$^{+/fl}$ *Cag*-CreER (Cag-CreER line: Jackson 04453), were injected intraperitoneally with Tamoxifen (Sigma T5648) (100 mg/kg body weight, 10 mg/ml stock solution in corn oil) or vehicle (corn oil). Animals, aged P30-P40, were injected for five consecutive days and behavior tests were performed >21 days post (last) injection (animals age P55-P65) to ensure successful recombination, protein degradation and Tamoxifen (and its metabolites) clearance, as previously described [58]. Behavioral tests were performed as described above. After behavioral tests, animals were sacrificed, the brain dissected and the right hemisphere was used for tissue lysis and western to determine Cul3 protein levels and success of Tamoxifen induced deletion. As described previously [59], in 5%–20% of mice treated with Tamoxifen, induction appeared to fail altogether, and no changes on Cul3 protein levels could be observed. These mice, and their corresponding vehicle treated littermates, were excluded from analysis.

**Statistics**

Statistical analyses were performed using Microsoft ® Excel ® 2013, Origin Software (Origin Inc.) and GraphPad Prism 6/8. Shapiro–Wilk test was used to evaluate normal distribution, means and standard deviations of the data. Parametric data were analyzed for significance using unpaired two-tailed t-tests, 1-way or 2-way ANOVAs with Sidak's post-hoc test, using $*P < 0.05$, $**P < 0.01$, and $***P<0.001$ for significance, and presented as bar, box and whiskers, scatter dot plots and mean ± standard error of the mean (SEM), unless otherwise specified. Data sets with non-normal distributions were analyzed using the two-tailed Mann-Whitney U test. Adjustment for multiple comparisons were made using post-hoc tests. Cumulative probability plots of the amplitude and inter-event interval of synaptic currents were compared with the Kolmogorov-Smirnov two-sample test. Detailed statistics are presented in Supplementary Table 1. Experiments were replicated at least three times.

**Data availability:** The data that support the findings of this study are available from the corresponding author upon reasonable request.


46 McEvoy, J. D., Kossatz, U., Malek, N. & Singer, J. D. Constitutive turnover of cyclin E by Cul3 maintains quiescence. *Mol Cell Biol* **27**, 3651-3666, doi:10.1128/MCB.00720-06 (2007).

47 Tarlungeanu, D. C. *et al.* Impaired Amino Acid Transport at the Blood Brain Barrier Is a Cause of Autism Spectrum Disorder. *Cell* **167**, 1481-1494 e1418, doi:10.1016/j.cell.2016.11.013 (2016).

48 Deliu, E. *et al.* Haploinsufficiency of the intellectual disability gene SETD5 disturbs developmental gene expression and cognition. *Nat Neurosci* **21**, 1717-1727, doi:10.1038/s41593-018-0266-2 (2018).

49 Arbuckle, E. P., Smith, G. D., Gomez, M. C. & Lugo, J. N. Testing for odor discrimination and habituation in mice. *Journal of visualized experiments : JoVE*, e52615, doi:10.3791/52615 (2015).

50 Gstrein, T. *et al.* Mutations in Vps15 perturb neuronal migration in mice and are associated with neurodevelopmental disease in humans. *Nat Neurosci* **21**, 207-217, doi:10.1038/s41593-017-0053-5 (2018).

51 Schindelin, J. *et al.* Fiji: an open-source platform for biological-image analysis. *Nature methods* **9**, 676-682, doi:10.1038/nmeth.2019 (2012).

52 Huang da, W., Sherman, B. T. & Lempicki, R. A. Systematic and integrative analysis of large gene lists using DAVID bioinformatics resources. *Nature protocols* **4**, 44-57, doi:10.1038/nprot.2008.211 (2009).

53 Huang da, W., Sherman, B. T. & Lempicki, R. A. Bioinformatics enrichment tools: paths toward the comprehensive functional analysis of large gene lists. *Nucleic acids research* **37**, 1-13, doi:10.1093/nar/gkn923 (2009).

54 Tinevez, J. Y. *et al.* TrackMate: An open and extensible platform for single-particle tracking. *Methods* **115**, 80-90, doi:10.1016/j.ymeth.2016.09.016 (2017).

55 Schaffer, A. E. *et al.* Biallelic loss of human CTNNA2, encoding alphaN-catenin, leads to ARP2/3 complex overactivity and disordered cortical neuronal migration. *Nature genetics* **50**, 1093-1101, doi:10.1038/s41588-018-0166-0 (2018).

56 van der Walt, S. *et al.* scikit-image: image processing in Python. *PeerJ* **2**, e453, doi:10.7717/peerj.453 (2014).

57 Seginer, A., Schmidt, R., Leftin, A., Solomon, E. & Frydman, L. Referenceless reconstruction of spatiotemporally encoded imaging data: principles and applications to real-time MRI. *Magn Reson Med* **72**, 1687-1695, doi:10.1002/mrm.25084 (2014).

58 Jahn, H. M. *et al.* Refined protocols of tamoxifen injection for inducible DNA recombination in mouse astroglia. *Sci Rep* **8**, 5913, doi:10.1038/s41598-018-24085-9 (2018).

59 Guenthner, C. J., Miyamichi, K., Yang, H. H., Heller, H. C. & Luo, L. Permanent genetic access to transiently active neurons via TRAP: targeted recombination in active populations. *Neuron* **78**, 773-784, doi:10.1016/j.neuron.2013.03.025 (2013).
